# Neuronal dynamics of cerebellum and medial prefrontal cortex in adaptive motor timing

**DOI:** 10.1101/2023.11.23.568408

**Authors:** Zhong Ren, Xiaolu Wang, Milen Angelov, Chris I. De Zeeuw, Zhenyu Gao

## Abstract

Precise temporal control of sensorimotor coordination and adaptation is a fundamental basis of animal behavior. How different brain regions are involved in regulating the flexible temporal adaptation remains elusive. Here we investigated the neuronal dynamics of cerebellar interposed nucleus (IpN) and medial prefrontal cortex (mPFC) neurons during temporal adaptation between delay eyeblink conditioning (DEC) and trace eyeblink conditioning (TEC). When mice trained for either DEC or TEC and subsequently subjected to a new paradigm, their conditioned responses (CRs) adapted virtually instantaneously. Changes in the activity of the IpN neurons to CR timing were prominent during DEC-to-TEC adaptation, but less so during TEC-to-DEC adaptation. In contrast, mPFC neurons could rapidly alter their modulation patterns during both adaptation paradigms. Accordingly, silencing of mPFC blocked the adaptation of CR timing. These results illuminate how cerebral and cerebellar mechanisms may play differential roles during adaptive control of associative motor timing.

## Introduction

Efficient coordination of daily movements often requires establishing associations with specific sensory inputs. Timing plays a crucial role in optimizing associative sensorimotor behavior. Certain movement components must occur at precise moments following the sensory cue^1^. For instance, predator needs to launch itself with precise timing to be able to catch a prey. To enhance the temporal alignment between sensory perception and desired motor responses, brain circuits must adapt the temporal aspects of sensorimotor associations^2, 3, 4, 5^. Understanding how the brain learns and produces temporally precise movements while maintaining flexible reaction timing remains a grand challenge in neuroscience.

Pavlovian eyeblink conditioning (EBC) offers an ideal experimental model to explore the precise temporal regulation of associative sensorimotor behaviors^6, 7, 8, 9^. In EBC training, an innocuous conditioned sensory stimulus (CS) is paired with an aversive unconditioned stimulus (US), which elicits an unconditioned eyeblink response (UR). Over repetitive CS-US pairing, CS-induced conditioned response (CR) gradually develops, facilitating a well-timed transformation from sensory input to motor output^6, 8, 9, 10, 11, 12^. The timing of CR is critically dependent on the temporal relationship between CS and US. In delay eyeblink conditioning (DEC), where the US occurs at CS cessation promptly, the CR aligns with the late phase of the CS epoch^10, 11, 12^. In trace eyeblink conditioning (TEC), the US is presented after a stimulus-free interval following CS termination, leading to a CR within this interval^6, 7, 8^. Thus, EBC training with distinct CS-US temporal patterns allows an identical sensory input to prompt movements with differing timings.

The cerebellum is a crucial region for acquiring and expressing CRs during EBC^13, 14, 15, 16, 17^. Mossy fiber and climbing fiber pathways transmit CS-and US-related inputs, converging onto the Purkinje cells in the cerebellar cortex^10,14^. By repetitive pairing these two inputs with consistent interval, well-timed neural modulation can be established in the cerebellar interposed nucleus (IpN), ultimately driving eyelid closure^10, 14, 18, 19, 20, 21, 22^. Indeed, both CS- and US-related activities are observable in Purkinje cells and IpN neurons during both DEC and TEC^6, 9, 10, 23^. Stimulating IpN neurons triggers prompt eyelid closure, whereas suppressing the IpN outputs significantly hampers both CR acquisition and expression^8, 9, 11, 12, 21, 22, 24, 25, 26^. This underscores the causal relationship between IpN activity and CR performance. Nonetheless, it remains unclear whether, once CRs are established, the temporal features of these learned movements can still be adaptively modified, and if so, the potential involvement of specific brain regions.

The medial prefrontal cortex (mPFC) is a higher order association cortex crucial for adaptive cognitive control of sensorimotor tasks, and is therefore likely to contribute to the flexible regulation of sensorimotor timing^27, 28^. Specifically, the mPFC plays a permissive role in gating TEC, but not DEC, highlights its importance for the temporal features of learned movements^14, 16, 29, 30, 31, 32, 33, 34, 35, 36^. It has been suggested that mPFC neurons transmit CS-related signals through the pontine nuclei to the cerebellum and establish a short-term memory trace that permits the CS-US association during TEC^33, 35, 37, 38^. Hence, we hypothesize that both cerebellum and mPFC are required for flexible control of sensorimotor timing, yet with differential impacts.

To test this hypothesis, we investigated shared and distinct neural dynamics of the IpN and mPFC neurons during DEC and TEC. We focused specifically on understanding how mice adjust their CR timing in response to the altered CS-US interval. We found that mice exhibited almost instantaneous adjustments of CR onset timing when subjected to adaptation from DEC-to-TEC or TEC-to-DEC paradigms. Remarkably, the temporal adaptation of CR was reflected in IpN neuron activity during DEC-to-TEC switch, but less so during TEC-to-DEC adaptation. In contrast, mPFC neurons had consistent alterations of their task related and baseline activity patterns during the paradigm switches. The physiological findings were in accordance with our perturbation experiments in that suppressing mPFC activity hindered animals’ ability of swiftly adapt CR timing during both DEC-to-TEC and TEC-to-DEC transitions. In summary, our results provide new insights into the cerebral and cerebellar mechanisms that underlie adaptive temporal control of learned associative movements.

## Results

### Distinct CR kinematics in mice trained with DEC and TEC paradigms

We first trained two separate cohorts of mice, using either the TEC or DEC paradigm, for direct comparison of their behavioral and neuronal characteristics. Both TEC and DEC trials began with a 250 ms flashlight stimulus serving as the CS, paired with a 15-ms air-puff to the left eye as the US. In the DEC paradigm, the US followed the CS immediately, while a 250 ms interval was introduced between the CS offset and the US onset in the TEC paradigm (Fig. 1a, c). Our experimental design aimed to investigate how identical sensory inputs, presented at different CS-US intervals, lead to distinct temporal input-output features in behavior after sensorimotor learning.

**Figure 1:**
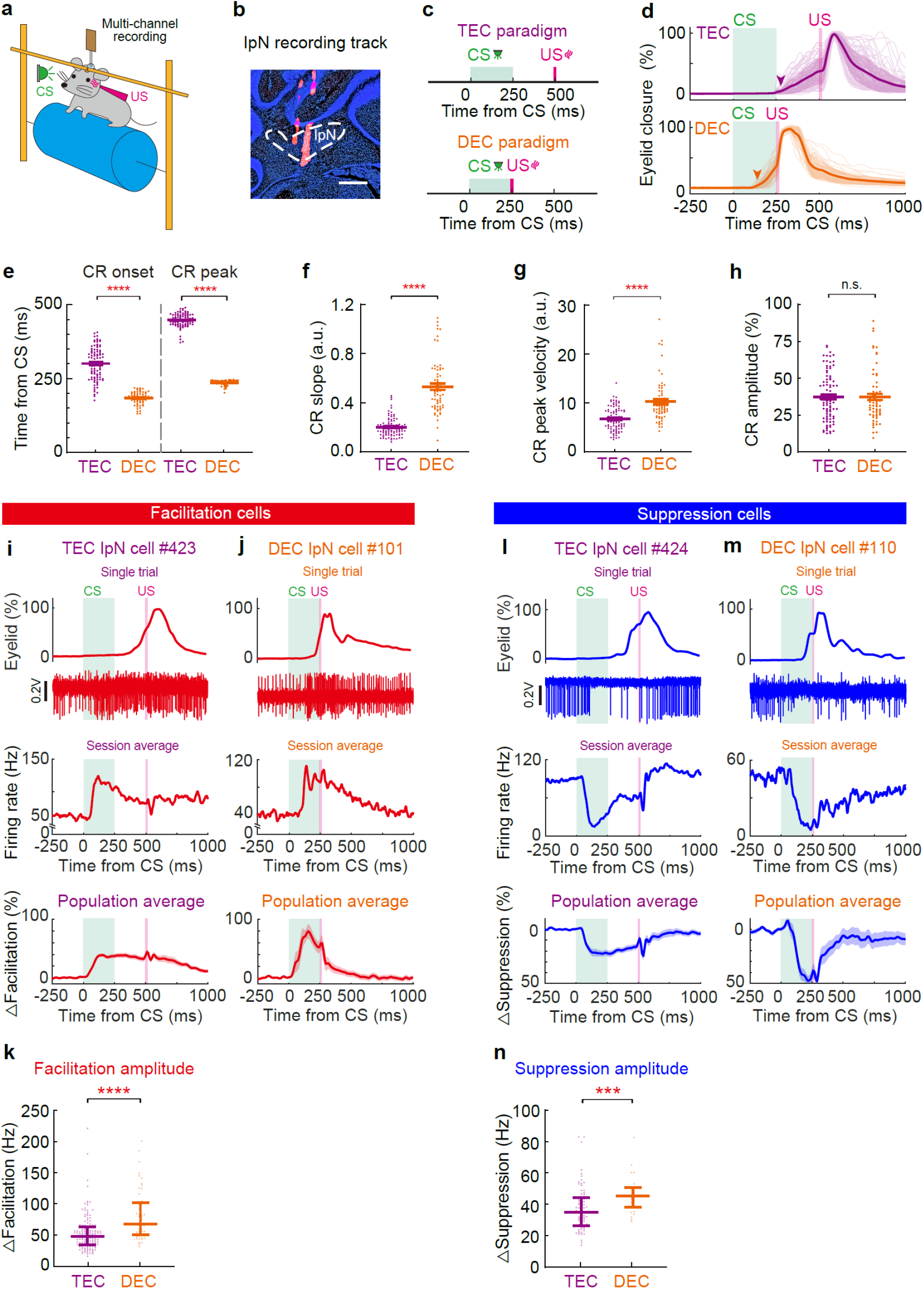
Mice establish well-timed eyelid closure in response to specific CS-US temporal relationships. **a** Schematic of the experimental setup. A head-fixed mouse is presented with a green light as the CS and an air puff as the US, with a multichannel silicon probe penetrating into the interposed nuclei (IpN). **b** An example of a DiI labeled recording track in the IpN (scale bar, 1 mm). **c** The CS-US relationships of the TEC and DEC paradigms. **d** Summary of eyelid traces from example TEC and DEC recording sessions. Arrowheads indicate the CR onsets. **e-h** Comparison of the CR onset and peak timings (**e**, *P* < 1.0 × 10^−15^ for both), CR slope (**f**, *P* < 1.0 × 10^−15^), CR peak velocity (**g**, *P* = 7.7 × 10^−10^), and CR amplitude (**h**, *P* = 0.85) between the TEC- and DEC-trained mice (*n* = 92 and 67 sessions). **i** Neuronal activities of IpN neurons in TEC-trained mice. Top, the eyelid closure curve and corresponding spike modulation from a single TEC trial. Middle, spike rate modulation of the same IpN neuron during a TEC recording session. Bottom, averaged firing activity of all IpN neurons with spike rate facilitation between CS-US intervals during TEC. **j** Same as (**i**) but for the DEC-trained group. **k** Comparison of the CR-related facilitation of IpN neurons in TEC- and DEC-trained mice (*P* = 4.4 × 10^−7^, *n* = 138 and 51 cells). **l-m** Same as **i-j**, but for the IpN neurons that showed CR-related suppression during TEC (**l**) or DEC (**m**). **n** Comparison of the CR-related suppression of IpN neurons in TEC- and DEC-trained mice (*P* = 0.0008, *n* = 71 and 27 cells). Data are shown as the mean ± s.e.m., except for (**d**), which is the mean ± SD, and for (**k**), (**n**), which are median with interquartile range. n.s.: not significant, ***: *P* ≤ 0.001, and ****: *P* ≤ 0.0001.

Mice developed distinct temporal features of their CRs following TEC and DEC training^8, 10, 11, 12^. TEC-trained mice had later CR onset and peak time compared to those of the DEC-trained mice (onset time: 300.38 ± 5.77 ms in TEC vs. 184.26 ± 2.56 ms in DEC, *P* < 1.0×10^−15^; peak time: 446.40 ± 2.30 ms in TEC vs. 235.88 ± 1.01 ms in DEC, *P* < 1.0×10^−15^; *n* = 92 and *n* = 67 sessions, Fig. 1d, e). However, the intervals between CR peak time to the US onset were comparable between TEC (27.37 ± 6.61 ms after US) and DEC (50.87 ± 7.02 ms after US) groups (*P* = 0.08, *n* = 92 and 27 sessions). Additionally, TEC-trained mice showed shallower CR slopes and lower CR peak velocities compared to the DEC-trained mice (CR slope: 0.20 ± 0.008 in TEC vs. 0.53 ± 0.026 in DEC, a.u., *P* < 1.0×10^−15^; CR peak velocity: 6.7 ± 0.25 in TEC vs. 10 ± 0.54 in DEC, a.u., *P* = 7.7×10^−10^, *n* = 92 and *n* = 67 sessions, Fig. 1d, f, g); whereas the CR amplitudes of TEC- and DEC-trained mice were comparable (40.23 ± 1.59% in TEC vs. 40.57 ± 2.00% in DEC, *P* = 0.85, *n* = 92 and 67 sessions, Fig. 1h). To rule out the possibility that distinct CR kinematics in TEC and DEC trained mice was influenced by specific detection method, we employed three well accepted methods to determine the precise CR onset timing^11, 39^. All three detection methods consistently illustrated the fundamental difference in CR timing in DEC and TEC training mice (Fig. 1e, Extended data Fig. 1a, b). These findings indicate that, despite the use of identical CS stimuli, the temporal specificity of the CS-US intervals determines specific CR timing and kinematics, which ensures the optimal conditioned eyelid closure before US onset.

We recorded the activity of cerebellar interposed nucleus (IpN) neurons ipsilateral to the trained eye during TEC and DEC (Fig. 1b). Among TEC-trained mice, a total of 138 IpN neurons exhibited CR-related facilitation (Fig. 1i, Extended data Fig. 1c), with a peak increase of 52.83 ± 2.48 Hz (Fig. 1k). For DEC-trained mice, we recorded 51 IpN neurons with facilitation, showing a peak increase of 80.71 ± 5.75 Hz (Fig. 1j, k, Extended data Fig. 1c). Additionally, we detected 71 suppression IpN neurons (with a suppression of 37.12 ± 1.75 Hz) from TEC-trained mice (Fig. 1l, n, Extended data Fig. 1d), and 27 IpN neurons (with a suppression of 45.92 ± 2.24 Hz) from DEC-trained mice (Fig. 1m, n, Extended data Fig. 1d). Despite the similarity in CR amplitudes between the two cohorts (Fig. 1h), the IpN modulation was notably larger in DEC-trained mice (Fig. 1k, n, facilitation groups: *P* = 4.4 × 10^−7^, n = 138 and 51 cells; suppression groups: *P* = 0.0008, n = 71 and 27 cells). These results unveil dynamic task-related neuronal activities within IpN, possibly reflecting a connection between the strength of IpN neuronal modulation and the CR kinematics.

### Temporal relationship between IpN activity and CR

CS-triggered IpN facilitation activates the downstream premotor nuclei, which consequently drives CRs in both TEC and DEC^11, 14, 16, 19, 40^. To what extent the temporal patterns of IpN neurons encode the temporal feature of CRs in DEC and TEC paradigms? We analyzed the temporal relationship between IpN modulation and CRs in DEC- and TEC-trained mice (Fig. 2a). The facilitation onset of IpN neurons consistently preceded CR onset in both TEC and DEC groups (Fig. 2b scatter plot, TEC facilitation onset vs. CR onset: 138.18 ± 8.29 ms vs. 295.59 ± 4.65 ms, *P* < 1.0 x 10^−15^, *n* = 138 cells. DEC facilitation onset vs. CR onset: 105.10 ± 5.84 ms vs. 189.60 ± 2.92 ms, *P* = 2.0 × 10^−15^, *n* = 51 cells). Despite the evident disparity in CR timings (Fig. 2b, right distribution histogram, *P* < 1.0 × 10^−15^), the onset timings of IpN modulation remained comparable between the TEC and DEC groups (Fig. 2b, top, *P* = 0.34). Consequently, a larger interval (ΔOnset) between IpN modulation and CR onset was observed in the TEC group (TEC Δonset: 157.41 ± 9.77 ms, DEC Δonset: 84.50 ± 6.40 ms, *P* = 1.58 × 10^−9^). To evaluate if IpN neurons encoded CR onset timing on a trial-by-trial basis, we conducted a temporal correlation analysis between modulation onset and CR onset for all CR-related IpN neurons. For a subgroup of IpN neurons from TEC-trained mice, their CR onset consistently aligned with the modulation onset on a trial-by-trial basis (Fig 2c, e correlation r = 0.72, *P* = 8.72×10^−6^ for the example neuron, *n* = 19). Similar correlations were also found in the DEC-trained mice (Fig 2d, f, correlation *r* = 0.77, *P* = 1.04 × 10^−5^ for the example cell, *n* = 7). Hence, a subset of IpN neurons precisely predict CR onset timing for both TEC and DEC mice.

**Figure 2:**
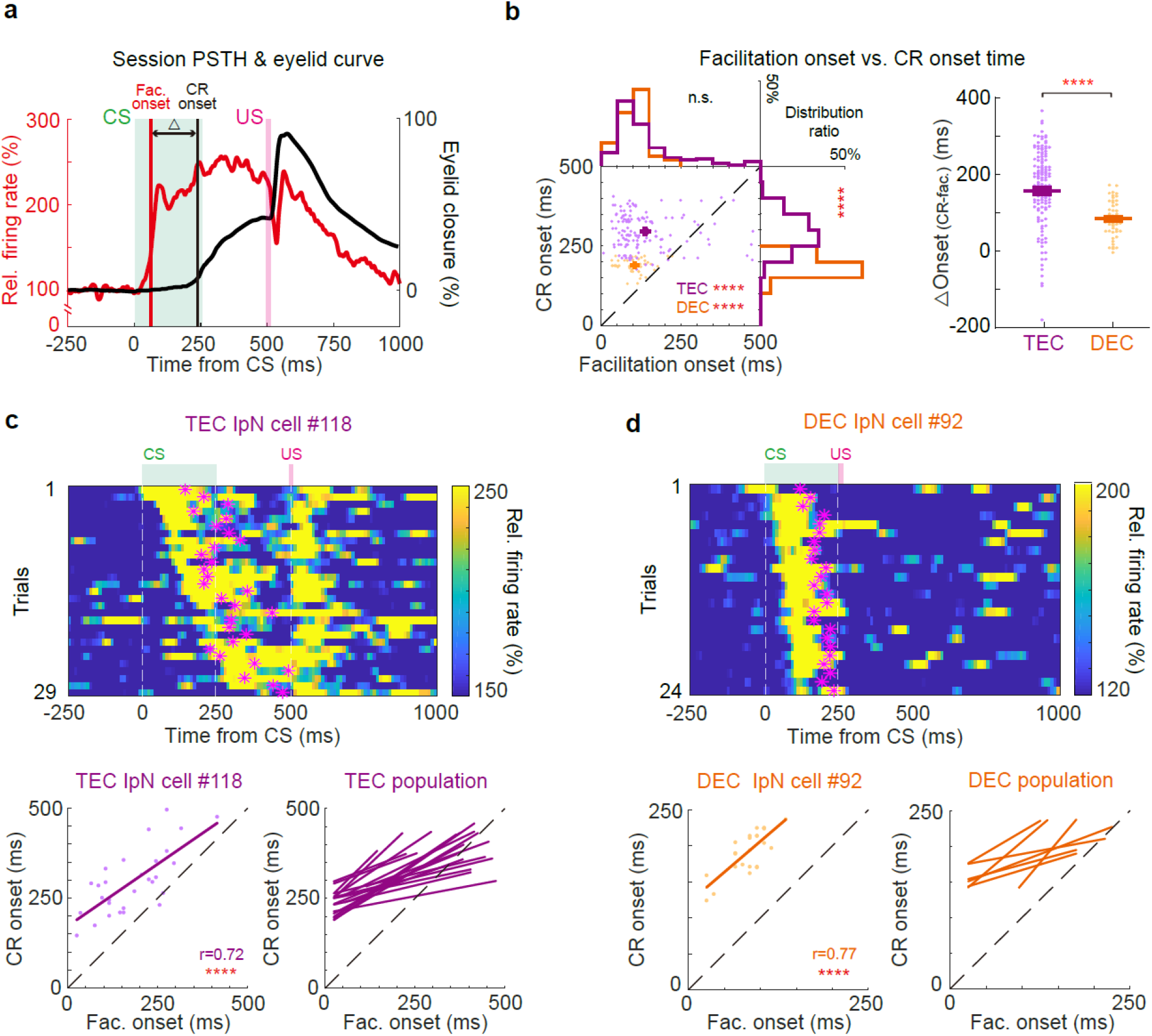
Weak temporal correlation between IpN population activity and CRs. **a** Illustration of the timing parameters extracted from the IpN neuron spike rate (red) and the eyelid closure (black) traces, including the timing of facilitation onset (Fac. onset) and CR onset, as well as their timing interval (△) in between. **b** Left, distribution and histograms of the IpN facilitation onset timing and the CR onset timing from TEC (purple) and DEC (orange) recordings. For both the TEC and DEC groups, IpN facilitation occurred prior to CR onset (TEC facilitation vs. CR onset, *P* < 1.0 × 10^−15^, DEC facilitation vs. CR onset, *P* = 2.0 × 10^−15^). The CR onset distribution in the DEC group was earlier than that in the TEC group (*P* < 1.0 × 10^−15^), whereas the distribution of IpN facilitation onsets was comparable in these two groups (*P* = 0.34, *n* = 138 and 51 cells). Right, the facilitation-to-CR onset interval (△) between TEC and DEC groups (*P* = 1.58 × 10^−9^, *n* = 138 and 51 cells). **c** IpN cells with significant trial-by-trial correlation between spike modulation onset and CR onset in TEC recordings. Top, heatmap of the instantaneous firing rate from an example IpN neuron during TEC trials, ordered by facilitation onset timing. CR onsets indicated by magenta stars. Bottom left, facilitation onset-CR onset pairs and their correlation during TEC trials from the example neuron (*r* = 0.72, *P* = 8.72 × 10^−6^). Bottom right, correlations of all TEC cells (*n* = 19) with this pattern. **d** Similar to **c** but for all the facilitation cells (*n* = 7) from the DEC group (*r* = 0.77, *P* = 1.04 × 10^−5^). Data are shown as the mean ± s.e.m., n.s.: not significant, ****: *P* ≤ 0.0001.

Previous studies have defined the ‘eyeblink-related IpN neurons’ as the neurons whose spike rates correlate with the CR amplitudes on a trial-by-trial basis^11, 12^. We then compared the timing of CR-related modulation in these eyeblink-related IpN neurons. Consistent with the findings from the population of all IpN neurons, the modulation onset of eyeblink-related IpN neurons had no significant difference between DEC- and TEC-trained mice, resulting in a significantly larger interval between IpN modulation and CR onset in TEC-trained mice (Extended data Fig. 2a-c). In addition, similar neuronal activity-behavior relationships were observed in the IpN neurons showing CR-related suppression (Extended data Fig. 2d). These results showed that the temporal relationships between IpN output and CR onset were differentially set for both DEC- and TEC-trained mice.

IpN output can directly initiate eyelid closure through a di-synaptic pathway involving the facial nucleus via the red nucleus^12^. We asked whether the interval between IpN modulation and CR onset is determined by the synaptic delay of this pathway. To assess this, we conducted direct electrostimulation of IpN neurons and measured the minimum induction time required to elicit eyelid closure via the IpN-red nucleus-facial nucleus pathway. With varying current strengths, we observed an increment in eyelid closure amplitudes while maintaining consistent eyelid closure onset timings (Extended data Fig. 3a). Remarkably, this minimum induction time was significantly shorter than the interval between IpN modulation and CR onset in both DEC- and TEC-trained mice (Extended data Fig. 3b-d). Additionally, IpN neurons showed clear CR related modulation during the trials in which mice did not show CR, although much less pronounced than the trials with CR (Extended data Fig. 3d, e). Hence, IpN modulation does not exhibit a fixed temporal relationship with CR onset that solely reflects the motor drive for eyelid closure. Nor does IpN modulation encode CR in an all or nothing manner. Instead, our data suggests the implementation of multiplex encoding strategies for controlling CR onsets in IpN.

We asked what types of encoding strategies IpN neurons could potentially implement to track the CR onset timing. We trained a decoding classifier to distinguish the early- and late-CR onset trials using the firing dynamics of IpN neurons recorded during TEC^41^. We sorted all the CR trials based on their onset timing and divided them into early- and late-onset trials. Subsequently, a subset of trials from both datasets were selected for training the decoder and the rest for assessing the decoding accuracy (Extended data Fig. 4a; see Methods). The decoding classifier showed that the CR onsets could be decoded based on the modulation timing in a subgroup of IpN neurons (Extended data Fig. 4b, *n* = 11 cells). Moreover, a distinct subset of IpN neurons facilitated their firing rates only during the late-CR trials (Extended data Fig. 4c, *n* = 22 cells), while another subgroup of neurons increased firing rates for the early-CR trials (Extended data Fig. 4d, *n* = 9 cells). We also attempted to decode the neuron activity for the CR onsets in DEC-trained mice. However, the decoding results were insufficient due to the minimal variety of CR onset timings during the DEC. Together, our analysis utilizing the decoding classifier confirmed that IpN neurons employ multiplex encoding strategies to regulate the temporal aspects of CRs. Both temporal and spike rate coding mechanisms can play roles in the trial-by-trial control of CR timing.

### Rapid adaptation of CR onset timing during the DEC-to-TEC adaptation paradigm

Once well-trained mice establish a specific CR onset timing, could they still adjust their CR timing when exposed to a novel CS-US interval? We introduced the TEC paradigm to the mice previously trained in DEC (Fig. 3a). Mice retained their CR percentages but had significantly smaller CR amplitudes after we switched to the TEC paradigm (Fig. 3b-d, Extended data Fig. 5a, b). Remarkably, animals rapidly adapted their CR timing to match the novel CS-US interval. Both CR onset and peak timings were significantly delayed following the DEC-to-TEC adaptation (Fig. 3b-g, Extended data Fig. 5c, session averaged CR onset in Fig.3g. DEC: 168.03 ± 4.28 ms, TEC: 197.47 ± 5.64 ms, *P* = 5.0 × 10^−5^, *n* = 38 sessions). The adaptation of CR timing occurred almost instantaneously upon exposure to the novel TEC paradigm. This rapid behavioral adaptation was consistently observed across all animals (Fig. 3c, e, f, comparing all 4 quarters of TEC epochs with the DEC epoch, all *P* < 0.001, *n* = 38 sessions). Notably the adapted CR timing still differed from that of TEC-trained animals, despite comparable CR amplitudes (Fig. 3g, TEC-trained CR onset: 241.44 ± 7.61 ms, *P* = 0.0002, *n* = 92 sessions, Extended data Fig. 5b, c).

**Figure 3:**
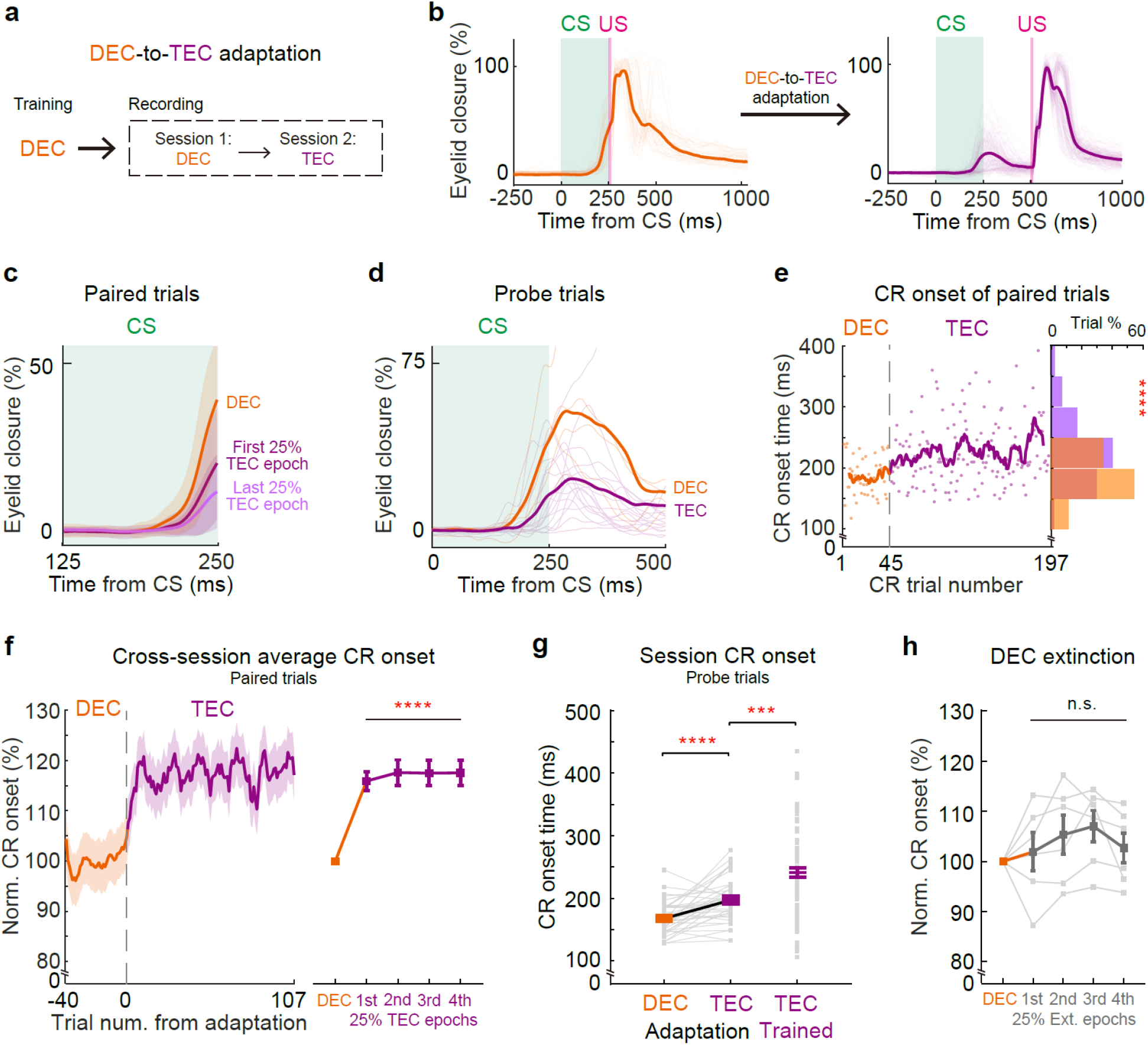
Rapid adaptation of CR onset timing during the DEC-to-TEC adaptation paradigm. **a** Experimental procedure for the animal training and paradigm switch to TEC in DEC-trained mice. **b** The eyelid closure curves of an example recording in DEC session (left) and TEC session (right). **c** The average eyelid closure curves from the paired trials of the example recording in **b** which is divided into DEC epoch (orange), first 25% TEC epoch (purple), and last 25% TEC epoch (light purple). **d** The average and individual (in dark and light colors) probe trial eyelid closure curves in the DEC (orange) and TEC (purple) sessions of the same recording. **e** The CR onsets of each trial during the example recording from DEC to TEC indicating rapid adjustment. The curve illustrates the moving average. The distribution histogram of the CR onsets in DEC (orange) and TEC (purple) is shown on the right (CR onset time DEC vs. TEC *P* = 4.54 × 10^−6^, *n* = 44 and 153 trials). **f** Left, cross-session averages of normalized CR onset timing from all DEC-to-TEC adaptation recordings. Right, statistical analysis of the CR onset timing for the DEC epochs and all four quartiles of TEC adaptation epochs (*P* < 0.0001 in all four comparisons, *n* = 38 sessions from 11 mice). **g** Comparisons of the CR onset timing of all DEC-to-TEC adaptation recordings together with that of the TEC-trained mice (*P* = 5.0 × 10^−5^ and 0.0002, *n* = 38 and 92 sessions). **h** The normalized CR onset timing from DEC extinction recordings, which are divided into DEC epochs and all four quartiles of extinction epochs (*P* = 0.23, *n* = 6 sessions). Data are shown as the mean ± s.e.m., n.s.: not significant, ***: *P* ≤ 0.001, and ****: *P* ≤ 0.0001.

Could the observed shift in CR onset timing during DEC-TEC training be attributed to a gradual extinction of the DEC response, paired with a simultaneous gradual acquisition of the TEC response? Using three distinct criteria to determine CR onset, we found that changes in CR amplitude had minimal impact on the timing of CR onset (see Extended Data Fig. 5d, e). To investigate this further, we trained another two separate cohorts of mice. One cohort underwent DEC training followed by extinction sessions. During extinction, these mice exhibited a gradual reduction in CR amplitude; yet, there was no corresponding delay in either CR onset or CR peak timing (Fig. 3h, *P* = 0.23, n = 6 sessions, Extended Data Fig. 6). In the second cohort, DEC-trained mice underwent TEC adaptation with an extended CS-US interstimulus interval, increased from 250 to 500 ms (i.e., DEC-to-TEC750, Extended Data Fig. 7a). Contrary to the possibility that CR could emergence with a much later timing, the shift in CR onset timing within this paradigm was rapid and comparable to that seen in the standard DEC-TEC paradigm (Extended Data Fig. 7b, c). These findings collectively indicate that mice can rapidly adjust CR onset timing, a mechanism that appears distinct from the extinction-and-acquisition model previously proposed in studies using rabbits^6^.

How do the cerebellar neurons response to such altered CS-US interval? One possibility is that IpN neurons alter their modulation patterns to generate adaptive CRs during the DEC-to-TEC adaptation. We focused on IpN neurons exhibiting CR-related facilitation in both DEC and TEC epochs (*n* = 40). Indeed, we observed adaptation in CR-related modulation upon the paradigm switch (Fig. 4a-d). On average the facilitation amplitude was decreased by 55.3%, and facilitation onset was delayed by 21.5% after the DEC-to-TEC adaptation (Fig. 4b-d, facilitation amplitudes: 164.64 ± 31.46% in DEC and 106.00 ± 19.10% in TEC, *P* = 0.0003; facilitation onset: 92.75 ± 7.13 ms in DEC, and 112.70 ± 9.33 ms in TEC, *P* = 0.001). However, the delayed facilitation onset didn’t perfectly align with behavioral changes. Adaptation in IpN modulation onset was significantly smaller than the adaptation of CR onset timing, leading to an extended latency between IpN neuronal activity and behavior (△Onset) after adaptation (Fig. 4e, f, DEC epoch: 70.16 ± 7.24 ms, TEC: 81.80 ± 8.80 ms, *P* = 0.0053). Similar phenomena were also observed in IpN neurons exhibiting CR-related suppression (Extended data Fig. 8). In summary, our data underscores that mice can rapidly recalibrate their CR timing in response to a novel CS-US interval. However, these changes in behavior cannot be solely accounted for by the adaptation of neuronal activity in IpN.

**Figure 4:**
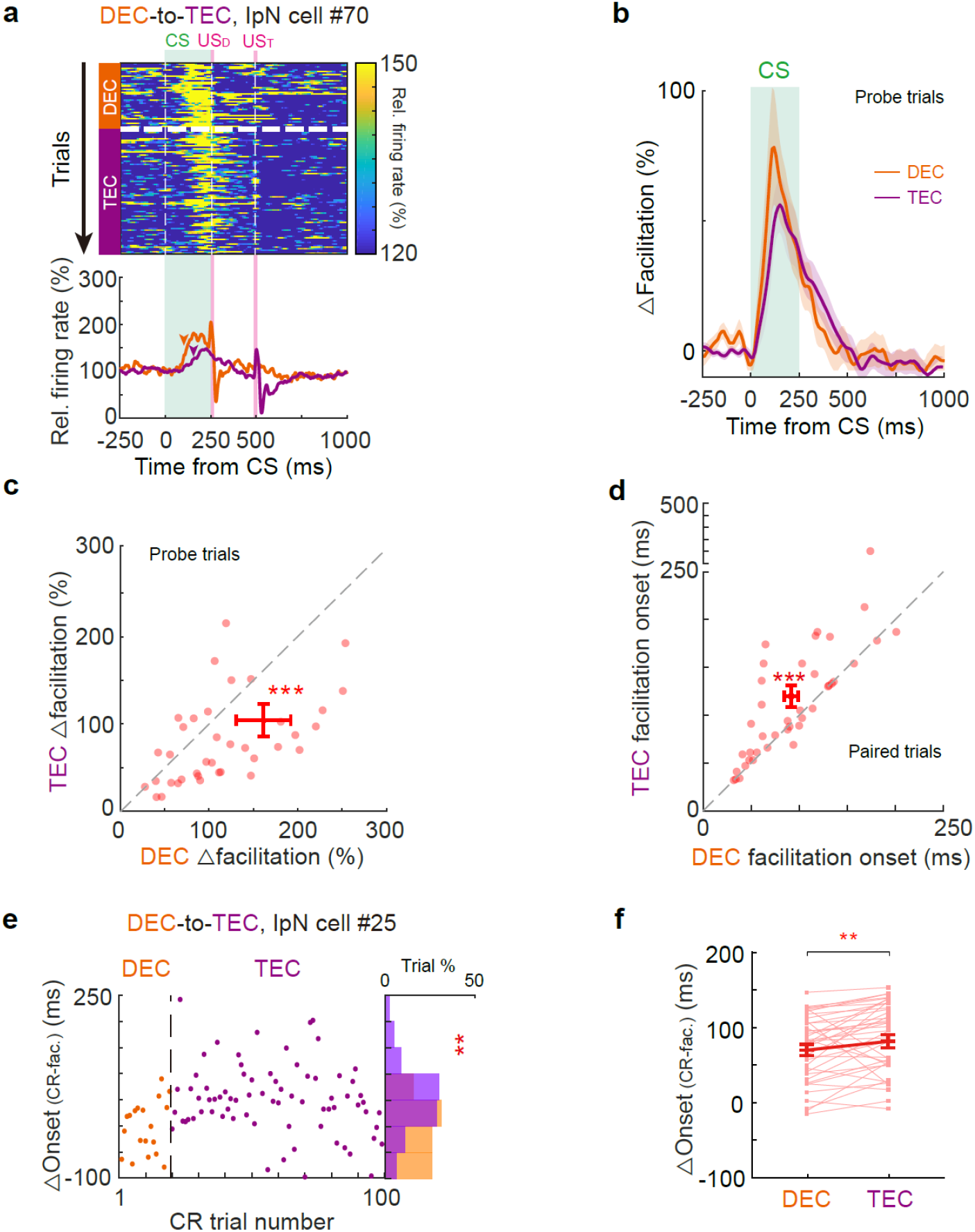
Adaptation of CR timing is not fully explained by the changes of IpN modulation during DEC-to-TEC adaptation. **a** Heatmap of trial-by-trial instantaneous firing rate (top) and the average firing rates (bottom) of an example IpN neuron during DEC-to-TEC adaptation. Arrowheads indicate the modulation onsets, US_D_ and US_T_ indicate the USs of DEC and TEC sessions. **b** Average normalized firing rates of all facilitated IpN cells (*n* = 40 cells from 27 sessions) during the recordings. **c** Pairwise comparison of the facilitation amplitude during the DEC and TEC sessions for all IpN facilitation neurons (*P* = 0.0003, *n* = 40 cells). **d** Same as (**c**), but for the facilitation onset timing (*P* = 0.001, *n* = 40 cells). **e** Left, the trial-by-trial interval (△) between the onset timings of CR and neuronal modulation from an example IpN cell during DEC-to-TEC adaptation. The distribution histogram of the △ value is shown on the right (*P* = 0.0097). **f** Summary of all facilitation IpN neurons indicating an increase in the average △ value after DEC-to-TEC adaptation (*P* = 0.0053, *n* = 40 cells). Data are shown as the mean ± s.e.m., **: *P* ≤ 0.01, and ***: *P* ≤ 0.001.

### Rapid adaptation of CR onset timing during the TEC-to-DEC adaptation paradigm

We proceeded to investigate whether the TEC-trained mice could flexibly adapt their CR timing when exposed with a shorter CS-US interval in a TEC-to-DEC adaptation (Fig. 5a). Intriguingly, TEC-trained mice also showed an adaptive response to this paradigm switch, displaying a shortened CR onset timing post-switch (Fig. 5b-g, in Fig.5g, TEC epoch: 232.30 ± 9.33 ms, DEC epoch: 173.82 ± 4.43 ms, *P* = 2.05 × 10^−8^, *n* = 33 sessions). Mirroring the outcome of the DEC-to-TEC adaptation, the TEC-to-DEC adaptation led to nearly instantaneous adjustments in CR onset timing. Within merely 10 trials post-adaptation, CR onsets were significantly shortened, with CR kinematics continuing to adapt throughout the novel DEC trials (Fig. 5c-f, comparing the 4 quartiles of the DEC trials with the TEC trials, all *P* < 0.001, *n* = 33 sessions). Decreases in CR percentage, increases in CR amplitude, and earlier CR peak time were observed during the TEC-to-DEC adaptation (Extended data Fig. 9a-c). However these changes in CR kinematics had no impact on CR onset time detection (Extended data Fig. 9d, e). By the end of the TEC-to-DEC adaptation, CR onset timing and amplitudes became indistinguishable from those mice trained with the DEC paradigm (Fig. 5g, DEC-trained CR onset: 168.02 ± 4.28 ms, *P* = 0.14, *n* = 38 sessions; Extended data Fig. 9b, c). Therefore, mice were capable of fully adapting the CR onset timing within a single TEC-to-DEC session.

**Figure 5:**
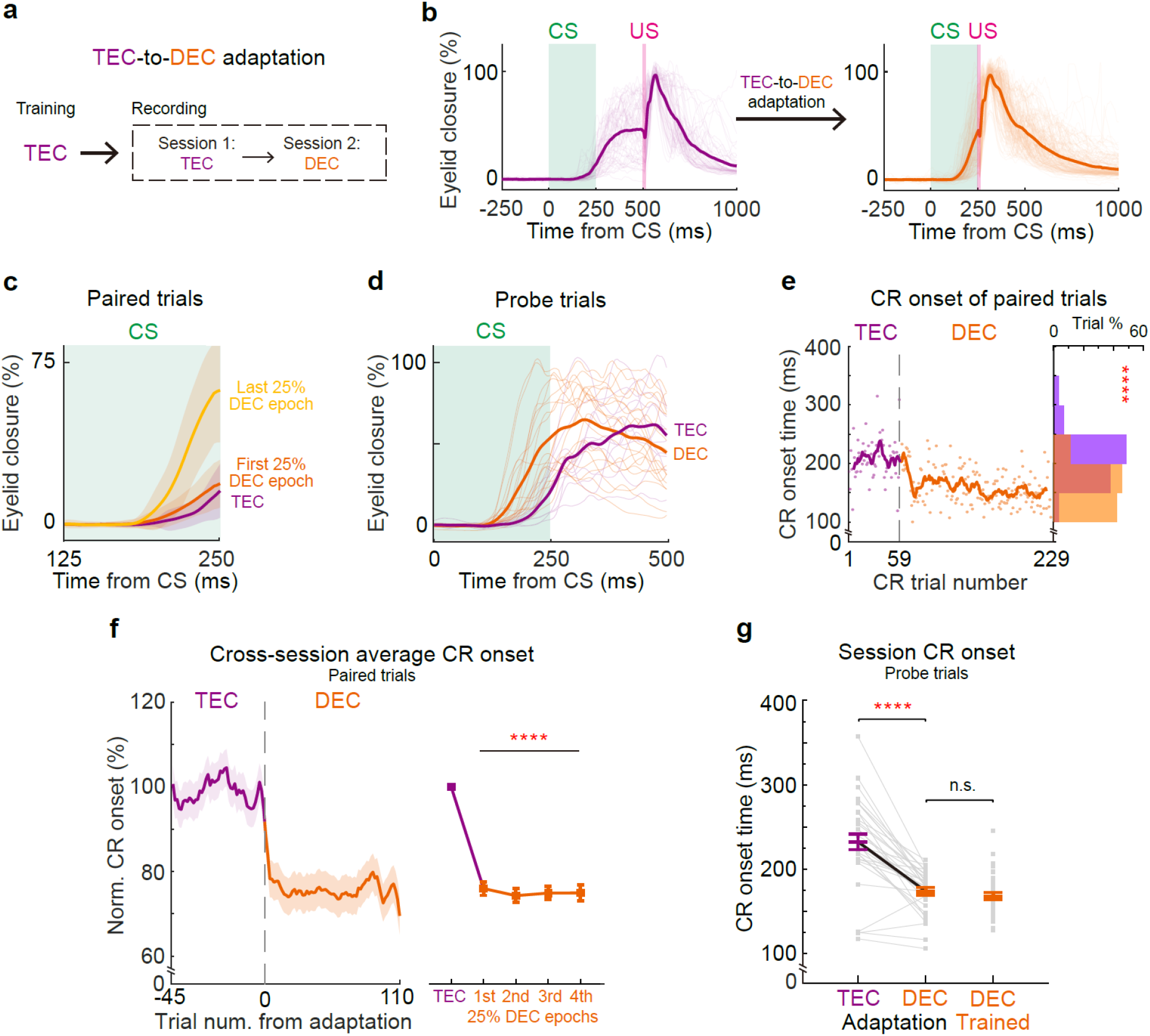
Rapid adaptation of CR onset timing during the TEC-to-DEC adaptation paradigm. **a** Experimental procedure for the animal training and paradigm switch to DEC in TEC-trained mice. **b** The eyelid closure curves of an example recording in the TEC (left) and DEC (right) sessions of TEC-to-DEC adaptation. **c** The average eyelid closure curves from the paired trials of the example recording in **b** which is divided into TEC epoch (purple), first 25% DEC epoch (orange), and last 25% DEC epoch (light orange). **d** The average and individual (in dark and light colors) probe trial eyelid closure curves in the TEC (purple) and DEC (orange) sessions of the same example session. **e** The CR onsets of each trial during the example recording from TEC to DEC indicating rapid adjustment. The curve illustrates the moving average. The distribution histogram of the CR onsets in TEC (purple) and DEC (orange) is shown on the right. (CR onset time TEC vs. DEC *P* < 1.0 × 10^−15^, *n* = 58 and 171 trials). **f** Left, cross-session averages of normalized CR onset timing from all TEC-to-DEC adaptation experiments. Right, statistical analysis of the CR onset timing for the TEC sessions and all 4 quartiles of DEC sessions (*P* < 1.0 × 10^−15^ in all four comparisons, *n* = 33 sessions from 14 mice). **g** Comparisons of the CR onset timing of all TEC-to-DEC adaptation experiments together with that of the DEC-trained mice (*P* = 2.05 × 10^−8^ and 0.14, *n* = 33 and 38 sessions). Data are shown as the mean ± s.e.m., ***: *P* ≤ 0.001, and ****: *P* ≤ 0.0001.

Could these changes in behavior be reflected in the activity of IpN neurons? We recorded IpN neurons, focusing on those CR-related facilitation across the TEC-to-DEC adaptation. Despite the swift adaptation of CR timing, we observed no substantial alterations in the modulation patterns of IpN neurons. Both the onset and amplitude of CR-related facilitation persisted unchanged after the TEC-to-DEC adaptation (Fig. 6a, b). Furthermore, pairwise comparisons indicated no statistical differences in facilitation amplitude and onset timing during the adaptation (Fig. 6c, d; facilitation amplitude: 119.19 ± 17.88% in TEC and 96.33 ± 7.35% in DEC, *P* = 0.40; facilitation onset: 106.31 ± 13.39 ms in TEC and 111.03 ± 8.90 ms in DEC, *P* = 0.65, *n* = 35 cells). However, the altered CR onsets without concurrent changes in IpN activity resulted in an even shorter interval between neuronal activity and behavior (△Onset) during DEC epochs, as exemplified by a specific neuron (Fig. 6e). This change was consistently observed across a broader facilitation neuron population (Fig. 6f; TEC: 129.10 ± 14.22 ms, DEC: 64.56 ± 9.97 ms, *P* = 1.07 ×10^−5^, *n* = 35 cells). The neuronal activity of IpN neurons exhibiting CR-related suppression also remained unaltered during TEC-to-DEC adaptation (Extended data Fig. 10). Therefore, we did not find significant neuronal adaptation in IpN that could explain the rapid adaptative behavior.

**Figure 6:**
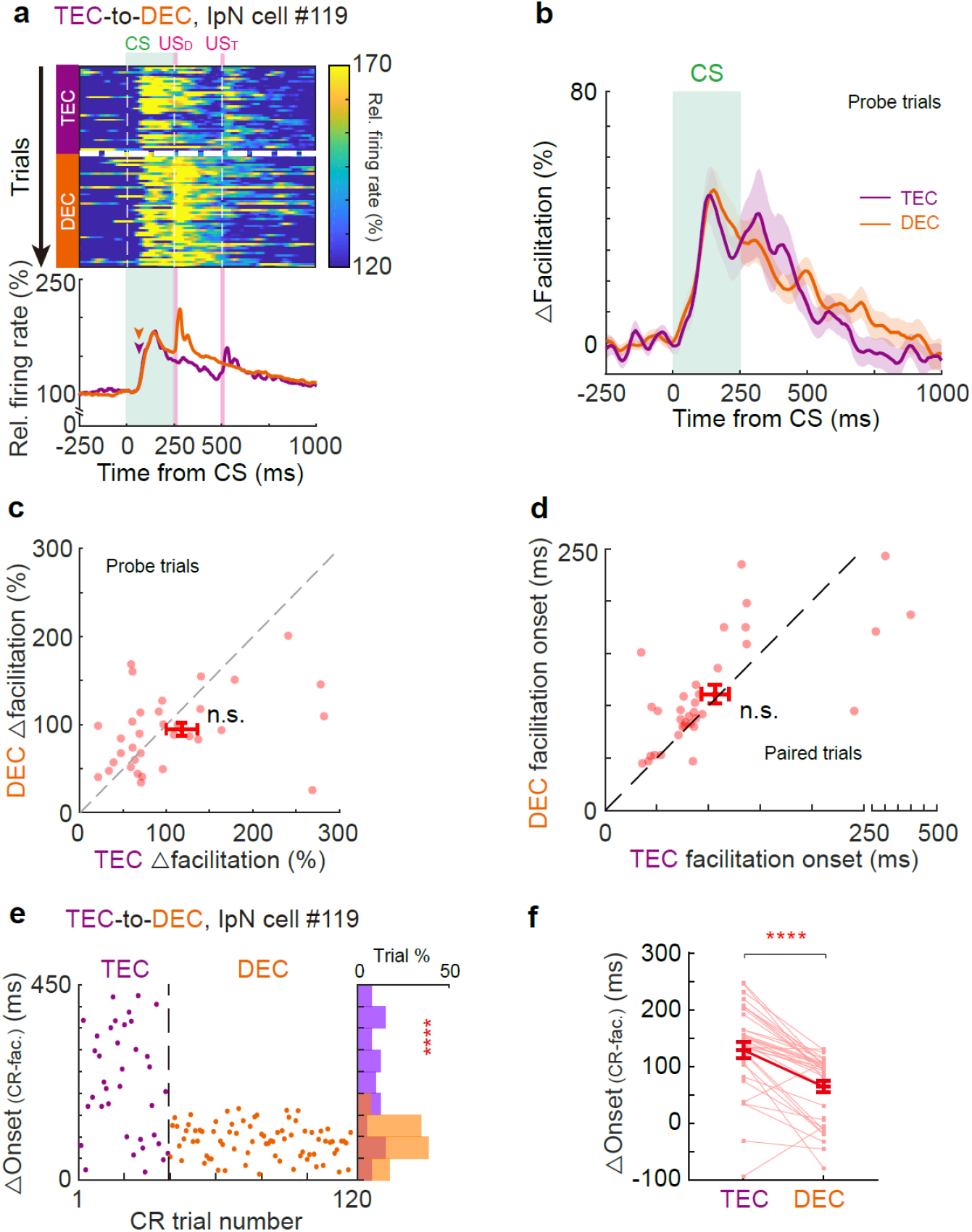
Adaptation of CR timing is not explained by the change of IpN modulation during TEC-to-DEC adaptation. **a** Heatmap of the trial-by-trial instantaneous firing rate (top) and average firing rates (bottom) of an example IpN neuron during the adaptation from TEC (purple) to DEC (orange). Arrowheads indicate the onsets of spike modulation, US_D_ and US_T_ indicate the USs of DEC and TEC sessions. **b** Average normalized firing rates of all facilitated IpN cells (n=35 cells from 25 sessions) during the TEC and DEC sessions. **c** Pairwise comparison of the facilitation amplitude during the TEC and DEC sessions for all IpN facilitation neurons (*P* = 0.40, *n* = 35 cells). **d** Similar to (**c**) but for the facilitation onset timing (*P* = 0.65, *n* = 35 cells). **e** Left, the trial-by-trial interval (△) between the CR and neuronal modulation onset timings from an example IpN cell during TEC-to-DEC adaptation. The distribution histogram of the △ value is shown on the right (*P* = 5.24×10^−6^). **f** Summary of all facilitation IpN neurons indicating a decrease in the average △ value after TEC-to-DEC adaptation (*P* = 1.07 × 10^−5^, *n* = 35 cells). Data are shown as the mean ± s.e.m., n.s.: not significant, ****: *P* ≤ 0.0001.

### CR-related activities in mPFC neurons

Our data demonstrates animals’ capacity to swiftly adapt their CR timings when introduced to novel CS-US intervals, nevertheless activity of IpN neurons did not fully correlate with this behavioral adaptation (Fig. 3-6). Moreover, the temporal adaptation occurred within a mere few trials, significantly faster than the time needed for associative learning that hinges on cerebellar long-term plasticity^8, 9, 12^. We posited that other brain regions might play a role in the rapid adaptation of CR timing. Previous studies have underscored the mPFC’s significance in TEC acquisition and expression^14, 16, 29, 30, 31, 32, 33, 34, 35, 36^. We sought to investigate whether the mPFC could be instrumental in orchestrating the bidirectional adaptation between TEC and DEC paradigms by recording the mPFC neurons located contralateral to the trained left eyes in mice during either TEC or DEC (Fig. 7a). Our recordings showed: 20.1% (64 out of 318) and 27.6% (94 out of 341) of mPFC neurons exhibited CR-related modulation in TEC- and DEC-trained mice, respectively (Fig. 7b, c, Extended data Fig. 11a, b). Among these neurons, a subset exhibited transient facilitation locked to the CS onset (Fig. 7d, f), with transient facilitation onsets of 111.63 ± 8.71 ms for the TEC group and 100.87 ± 4.67 ms for the DEC group. Several mPFC neurons displayed sustained modulation, and intriguingly, some of them exclusively modulated during CR trials, but not during non-CR trials (Fig. 7e, g, Extended data Fig. 11c, d). This distinct pattern suggests the mPFC activity is closely associated with conditioned eyelid closure during DEC and TEC.

**Figure 7:**
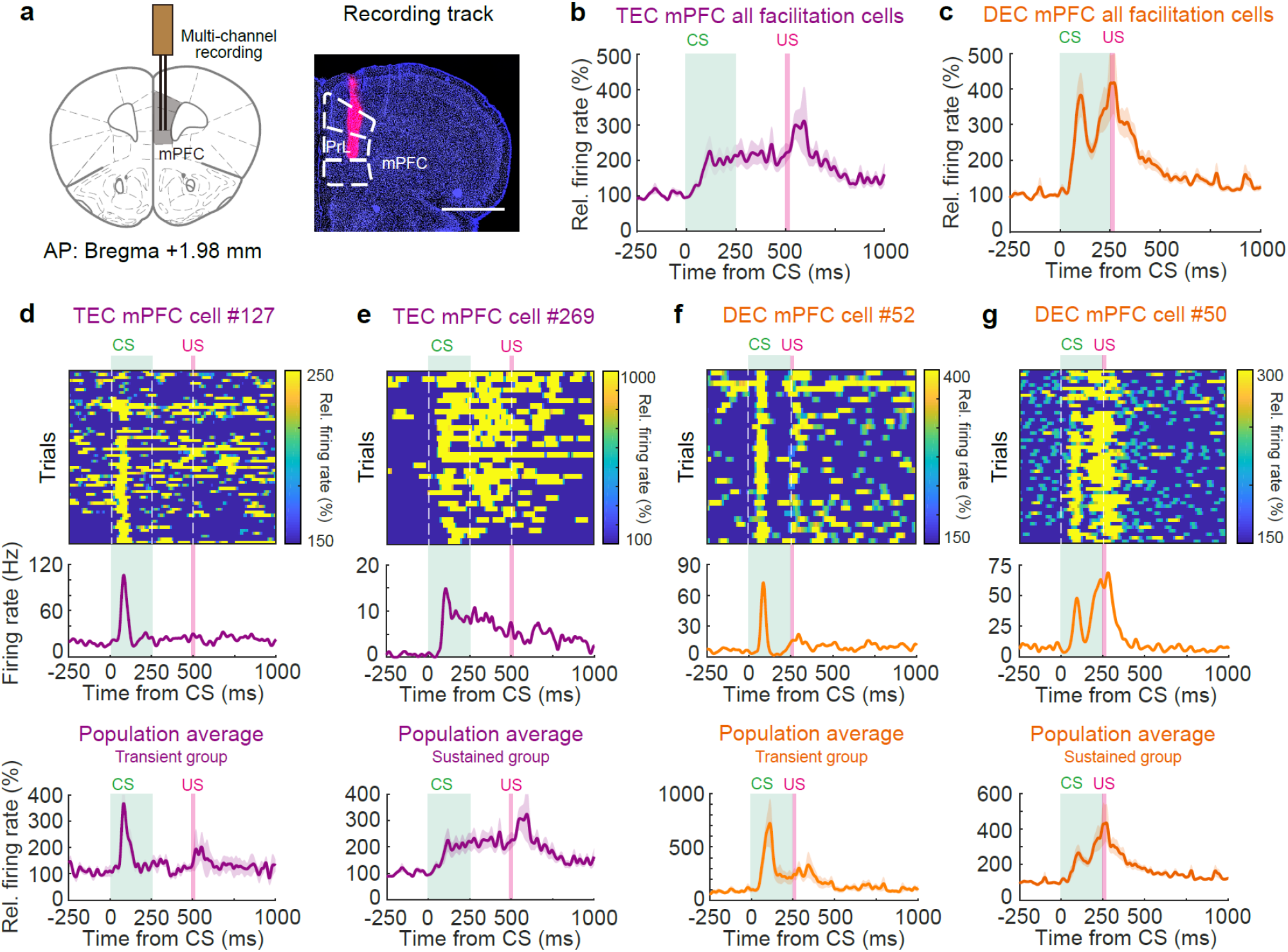
TEC- and DEC-related modulation observed in the mPFC. **a** Left, schematic of the mPFC recording site projected to the mouse brain atlas. Right, an example DiI-labeled recording track (PrL, prelimbic area, scale bar, 1 mm). **b** Average relative firing rates of all mPFC cells showing CR-related activities during TEC recordings (*n* = 64 cells, from 3 mice). **c** Similar to (**b**) but for the DEC recordings (*n* = 94 cells, from 3 mice). **d-g** Top and middle, the instantaneous firing rate heatmaps (top) and session average firing rate (middle) of example mPFC neurons, two cells during TEC and two cells during DEC. Bottom, population average of mPFC neuronal activity with specific modulation patterns during TEC and DEC recordings. **d-e** Neurons with transient (**d**) or sustained (**e**) facilitation recorded during the TEC paradigm (*n* = 5 and 59 cells). **f-g** Neurons with transient (**f**) or sustained (**g**) facilitation during the DEC paradigm (*n* = 18 and 78 cells).

### Rapid temporal adaptation of CR onset timing is encoded in mPFC neurons

We then investigated the involvement of mPFC neurons during the adaptation of the two paradigms, DEC-to-TEC and TEC-to-DEC, respectively (Fig. 8a, g). Consistent with the previous findings (Fig. 3f, 5f), both cohorts of mice exhibited rapid adaptation in response to the novel CS-US interval (Fig. 8b, h, *P* < 0.0001 for both, *n* = 11 DEC-to-TEC and *n* = 8 TEC-to-DEC sessions). Concurrent with these changes in CR onset, a subset of mPFC neurons displayed decreased modulation amplitudes during the DEC-to-TEC adaptation (Fig. 8c, d, *P* = 0.0003 for CR onset and *P* < 0.0001 for modulation, *n* = 65 DEC, and *n* = 71 TEC trials in Fig. 8c for the example recording, *P* = 0.0005, *n* = 29 cells in Fig. 8d for population). Interestingly, another subpopulation of mPFC neurons showed a higher baseline firing in response to the adaptation (Fig. 8e, f, *P* = 0.02 for CR onset and *P* < 0.0001 for baseline firing rate, *n* = 36 DEC, and *n* = 82 TEC trials in Fig. 8e for example recording, *P* < 0.0001, *n* = 133 cells in Fig. 8f for population). During TEC-to-DEC recordings, we also identified multiple mPFC neurons displaying either decreased modulation (Fig. 8i, j) or increased baseline firing rate (Fig. 8k, l) throughout the DEC epoch (*P* < 0.0001 for CR onset and *P* = 0.04 for modulation, *n* = 41 TEC, and *n* = 38 DEC trials in Fig. 8i for example recording, *P* = 0.0004, *n* = 12 cells in Fig. 8j for population; *P* < 0.0001 for both CR onset and baseline firing rate, *n* = 51 TEC, and *n* = 38 DEC trials in Fig. 8k for example recording, *P* < 0.0001, *n* = 75 cells in Fig. 8l for population). In summary, our results illustrate altered CR-related modulations and baseline activities in mPFC neurons during paradigm adaptation, implying that mPFC neurons could potentially signal the paradigm switch and guide the adaptive changes in CR timing.

**Figure 8:**
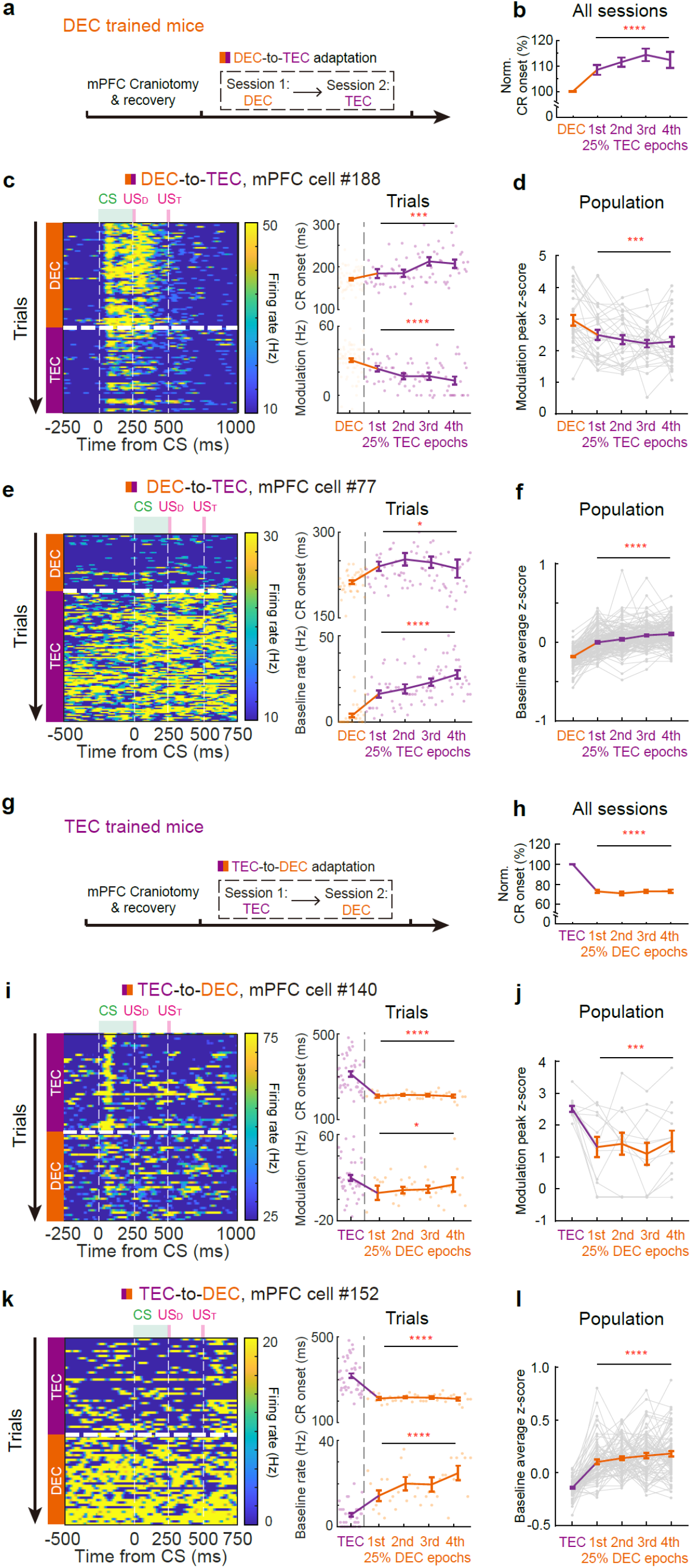
Adaptation of the CR-related modulations and baseline activities of mPFC neurons during paradigm switching. **a** Experimental procedures of DEC-to-TEC adaptation with mPFC recordings. **b** Statistical analysis of the CR onset timing for the DEC epochs and all four quartiles of TEC adaptation epochs with mPFC recording cohort (*P* < 0.0001, *n* = 11 sessions). **c** Left, firing activity heatmap of an example neuron with significantly reduced CR-related modulation during the TEC epoch of DEC-to-TEC adaptation. Right, the trial-by-trial CR onset (top, *P* = 0.0003) and the modulation (bottom, *P* < 0.0001) during the same recording session (*n* = 65 DEC, and *n* = 71 TEC trials). The changing trend is shown by the mean values of DEC epoch and four quartiles in TEC epoch. **d** The z-scored modulation peak during DEC epoch and four quartiles in TEC epoch with the mPFC neurons showing decreased modulation in TEC epochs of the adaptation (*P* = 0.0005, *n* = 29 cells). **e-f** Similar to (**c-d**), but for neurons with increased baseline firing rate during the TEC epoch of DEC-to-TEC adaptation (*P* = 0.02 for CR onset and *P* < 0.0001 for baseline firing rate, *n* = 36 DEC, and *n* =82 TEC trials in **e**, *P* < 0.0001, *n* = 133 cells in **f**). **g** Experimental procedures of TEC-to-DEC adaptation with mPFC recordings. **h** Similar with (**b**) but for TEC-to-DEC adaptation with mPFC recording cohort (*P* < 0.0001, *n* = 8 sessions). **i-j** Similar to (**c-d**), but for reduced modulation in the DEC epochs of TEC-to-DEC adaptation (*P* < 0.0001 for CR onset and *P* = 0.04 for modulation, *n* = 41 TEC, and *n* =38 DEC trials in **i**, *P* = 0.0004, *n* = 12 cells in **j**). **k-l** Similar to (**e-f**), but for neurons with increased baseline firing rate during the DEC epoch of TEC-to-DEC adaptation (*P* < 0.0001 for both CR onset and baseline firing rate, *n* = 51 TEC, and *n* =38 DEC trials in **k**, *P* < 0.0001, *n* = 75 cells in **l**).

### mPFC activity is essential for the temporal adaptation of CRs

To illustrate the functional relevance of mPFC in instructing the temporal adaptation, we injected GABA_A_ receptor agonist muscimol, in the mPFC of either DEC or TEC trained mice prior to the paradigm adaptation (Fig. 9a, b, Extended data Fig. 12a, b). With bilateral muscimol injections, we observed a reduction in CR percentage, although not complete elimination, in both DEC- and TEC-trained mice (Extended data Fig. 12c, d). Importantly, bilaterally inhibiting the mPFC was sufficient to abolish the rapid adaptation of CR onsets during both the DEC-to-TEC and TEC-to-DEC adaptations (Fig. 9c-f, in Fig. 9d, *P* = 0.97 and 0.054, respectively, *n* = 11 sessions, in Fig. 9f, *P* = 0.89 and 0.45, respectively, *n* = 15 sessions). These findings support the functional involvement of mPFC in regulating the temporal adaptation of CRs, suggesting that the frontal cortex and cerebellum might work in tandem to dynamically adjust the precise relationship between sensory input and motor output, allowing for the flexible modulation of behavior based on specific task requirements.

**Figure 9:**
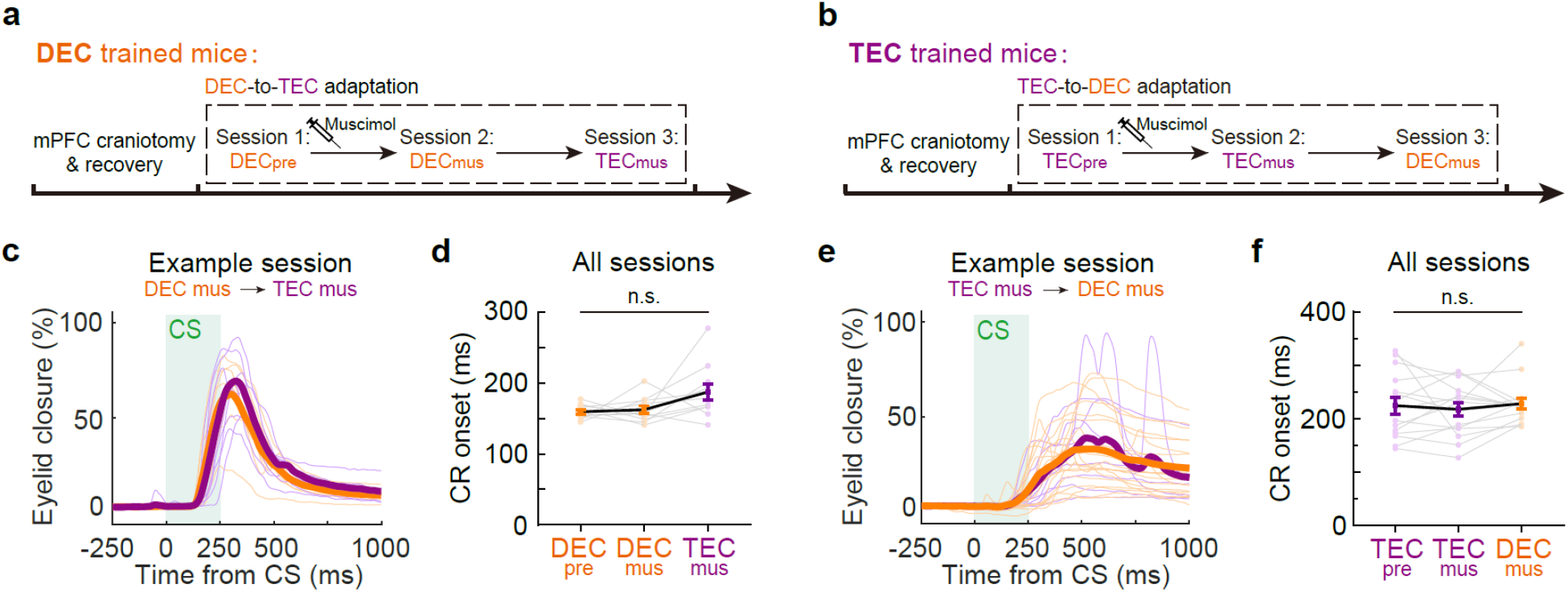
mPFC activity is essential for the adaptation of CR timing during paradigm switching. **a** Experimental procedures of mPFC inhibition during DEC-to-TEC adaptation. The recordings include three epochs: DEC prior to muscimol injection (DECpre), DEC after muscimol injection (DECmus) and the paradigm switch to TEC (TECmus). **b** Similar to (**a**) but for the TEC-to-DEC adaptation. **c** Eyelid closure curves of the CS-only trials during DEC (orange) and TEC (purple) sessions from an example session. **d** Summary of CR onset times for all mPFC inhibition during DEC-to-TEC sessions (*P* = 0.97 and 0.054, *n* = 11 sessions from 3 mice). **e** Same as (**c**), but for an example TEC-to-DEC session. **f** Same as (**d**) but for all TEC-to-DEC sessions (*P* = 0.89 and 0.45, *n* = 15 sessions from 3 mice). Data are shown as the mean ± s.e.m., n.s.: not significant.

## Discussion

The dynamic regulation of action timing plays a crucial role in guiding appropriate movements, yet the neural mechanisms regulating the temporal feature of sensorimotor behaviors remain largely elusive. In this study, we leveraged the well-understood neuron circuits underlying the sensory-cued eyelid closure response and examined the neuronal characteristics of cerebellar and mPFC circuits for the precise temporal regulation of conditioned responses. Focusing on animals trained with an identical CS but distinct CS-US intervals, our study reveals the common and distinct cerebellar coding features for DEC and TEC paradigms (Fig. 1,2). By examining the neural dynamics during DEC-to-TEC and TEC-to-DEC adaptations, we uncovered the characteristics of cerebellar activity underlying the rapid adaptation of action timing (Fig. 3-6). We further established the essential role of the mPFC in mediating such bidirectional adaptation (Fig. 7-9). All these findings support the role of cortico-cerebellar circuits for precise and adaptive temporal control of sensorimotor behaviors.

### Multimodal encoding of IpN activity during DEC and TEC

We have extensively examined the neuronal and behavioral dynamics in mice trained using either the DEC or TEC paradigm. The fundamental objective of Pavlovian eyeblink conditioning is to attain precise eyelid closure timing before the presentation of US. A wealth of studies has illuminated that this anticipatory response is facilitated by task-related modulation in IpN neurons, which occurs during the learning process^8, 9, 10, 11, 13, 14, 15, 16, 18, 19, 20, 21, 22, 24, 25, 26, 42, 43, 44, 45, 46^. Given that IpN neurons directly influence downstream premotor neurons in the red nucleus, it is postulated that CR-related modulation in IpN neurons is in close correlation with CR timing^9, 11, 24, 26^. Our comprehensive trial-by-trial and decoding analyses investigating the relationship between CR onset timing and IpN facilitation unequivocally validate the predictive role of IpN neurons. Notably, the timing of IpN facilitation precedes and forecasts the timing and kinematics of CR (Fig. 1-2). Consequently, our findings strongly endorse the perspective that IpN neurons play a pivotal role in the regulation of CR timing and kinematics within both the DEC and TEC paradigms.

Whether IpN neurons employ a common coding strategy to regulate CR timings in both DEC and TEC paradigms has not been systematically explored in mice. We conducted recordings of IpN neurons during both DEC and TEC paradigms utilizing high-density multiunit silicon probes. Mice trained in the DEC and TEC paradigms, which comprise an identical CS but different CS-US intervals, differ in their CR timings but not CR amplitudes (Fig. 1). As CR-related modulation in the IpN is considered the key driver of eyelid closure^9, 11^, one would assume that the IpN modulation in DEC- and TEC-trained mice also differs in timing but not in amplitude. Detailed activity-behavior correlation analysis revealed, however, that DEC-trained mice showed larger task-related modulation amplitudes. In contrast, the modulation onset timings at the population level were indistinguishable between TEC- and DEC-trained mice (Fig. 2), resulting in a much longer delay between IpN modulation and CR onset in TEC mice. Focusing on the ‘eyeblink neurons’, which were defined by their significant trial-by-trial activity-behavior correlation, we still observed clear difference in the activity-behavior intervals between TEC- and DEC-trained mice (Extended data Fig. 2). The delays between neural activity and behavioral response are an order of magnitude longer than the synaptic delays observed in CR onset following direct electrical stimulation of the IpN (Extended data Figure 3). These data suggest that information conveyed by the IpN is complex and cannot be viewed as merely a ‘motor command for eyelid closure’ and instead reflects a multimodal encoding pattern.

The complexity of these signals can stem from both cerebellar and extracerebellar processing^47, 48^. During EBC, CS- and US-related inputs converge at the cerebellar cortex, inducing persistent, input-specific synaptic depression at the parallel fiber to PC synapses^49^. The resultant suppression of simple spike activity in PCs, driven by long-term depression, is considered a central driver of CRs^6, 46^. Previous studies have suggested that the CR-related activities of PCs in animals transitioning from DEC to TEC paradigms share remarkable similarities and can be explained by a common inverse firing rate model^6^. However, more recent research challenges this notion, revealing that both facilitatory and suppressive modulations occur in PCs during DEC^50^. Consistently, we observed both task-related facilitation and suppression in the IpN neurons of both DEC- and TEC-trained mice. This challenges the oversimplified perspective that views IpN modulation solely as a motor command, favoring instead the notion of multiplex coding for precision and adaptability. Beyond the sensory domains, extracerebellar regions such as the hippocampus and mPFC have been implicated in contributing to TEC expression^8, 37, 51, 52, 53, 54^. The mossy fiber-granule cell pathway, which conveys these cognitive signals, theoretically follows the same plasticity rules as the sensory pathway, thereby providing additional task-related information to the IpN. A third stream of task-related information might stem from nucleo-cortical feedback loops, which have been proposed to offer predictive signals during DEC training^55, 56, 57^. These diverse input patterns might be beneficial for extending high-dimensional representations at the cerebellar input layer, thereby supporting behavioral state-specific computation in IpN^58^. Additionally, converging mossy fiber and climbing fiber collaterals onto the IpN could carry supplementary CS- and US-related information^11, 59^. This is substantiated by studies showing that mossy fiber collaterals undergo structural^59^ and synaptic plasticity^60^, thus directly shaping IpN neuron activities during behavior. In summary, these circuit mechanisms collectively augment the array of encoding dynamics in cerebellar output, leading to a richer and more nuanced signaling landscape.

### Cortico-cerebellar communication and rapid temporal adaptation

CR timing is determined by the specific temporal relationship between the CS and US during learning^61, 62^. Recent research indicates that cerebellar granule cells, optimized for encoding high-dimensional representations, form the foundation for temporal information encoding in sensorimotor behaviors^58^. Learning-triggered long-term plasticity at the synapses between parallel fibers and PCs is responsible for selecting relevant inputs, thereby shaping PC output patterns^46^. It might be expected that, in response to an abrupt change in CS-US interval, animals would temporarily maintain their CR onset timing, gradually adjusting to the new timing over training, following the time course of synaptic plasticity. Surprisingly, our findings counter this expectation, as mice trained under DEC or TEC paradigms exhibited the capacity to instantly adapt CR timings within a single training session upon the introduction of novel CS-US intervals (Fig. 3, 5). Such swift adaptation can manifest within just a few trials, significantly faster than the temporal dynamics of long-term synaptic plasticity in the cerebellum^63^. During this rapid adaptation, other cortico-cerebellar mechanisms are likely at play. In the conventional framework, CR timing in DEC primarily arises from cerebellar computation, while TEC engages an additional mPFC-cerebellar pathway that bridges the CS-free interval^33, 37^. The sequential activation of two sets of mossy fibers, encoding sensory and mPFC inputs, collectively shapes the CR’s temporal profile during TEC^62^. The latter set may be selectively disengaged, whereas the cerebellar computation of the sensory pathway prevails during TEC-to-DEC adaptation. To this end, cerebellar computation is expected to be similar in TEC-to-DEC converted mice and DEC-trained mice. This would find support from the results that the CR timing after TEC-to-DEC adaptation was identical to the CR timing of the DEC mice (Fig. 5g). Following the same reasoning, it is possible that novel extracerebellar information that emerges during DEC-to-TEC adaptation drives additional adaptation of IpN activity (Fig. 4) and delays the CR onset timing (Fig. 3). However, the DEC-to-TEC adaptation is unlikely to be fully dependent on the mPFC-cerebellar pathway. Mice trained with DEC were never exposed to the stimulus-free interval prior to adaptation; therefore, it is highly unlikely that they would activate the appropriate set of mossy fibers to bridge the CS-free interval^33, 37^. Yet, the switch in CR onset timing was as rapid as that seen in TEC-DEC adaptation. This stands in stark contrast to previous DEC-to-TEC studies in rabbits, where rabbits required prolonged training to gradually extinguish the original DEC response and reacquire a TEC response^6^. In line with our data using different TEC intervals (Extended Data Figure 7), we conclude that rodents possess the ability to rapidly adapt the timing of acquired CR responses within a short time frame without the need for extensive extinction and reacquisition of new CS-US intervals.

We show that the mPFC is critically involved in modulating CR expression as well as in facilitating rapid DEC-to-TEC and TEC-to-DEC adaptations (Fig. 7-9), consistent with the prominent roles of mPFC in TEC learning^14, 16, 29, 30, 31, 32, 33, 34, 35, 36^. This study sheds light on novel roles of the mPFC in both TEC and DEC paradigms. The phasic and persistent modulation patterns in the mPFC align with previous findings from TEC-trained rabbits^33, 36^, suggesting a multiplex coding of sensorimotor association in this region. This complexity arises from the coexistence of multiple components, including sensory, motor command, and outcome information^33, 36, 64^. Interestingly, prominent mPFC modulation time-locked to CS onset was observed not only in TEC-trained mice but also in DEC-trained mice. Furthermore, when we pharmacologically inhibited the mPFC, the suppression of CR percentage was observed in both DEC- and TEC-trained mice (Extended data Fig. 12). These results do not necessarily contradict the notion that the mPFC is involved in TEC but not DEC learning^29, 45, 65, 66, 67^. Although mPFC activity may not be necessary for DEC acquisition because the CS is directly contingent on the US, it could still influence animals’ internal states^68, 69, 70^, which has been shown to contribute to CR expression in DEC-trained animals^71, 72^.

Our findings also offer new insights into the mechanisms of temporal adaptation within the mPFC. The mPFC is known to play a significant role in various high-order cognitive functions, including working memory^73, 74, 75, 76, 77^, attention^78, 79^, and sensorimotor integration^80, 81^. Given the complexity of mPFC activity during cognitive control tasks, it is oversimplified to assume that the sole function of mPFC in eyeblink conditioning is to bridge the CS-US interval. It is therefore plausible that, in addition to the projections from the mPFC to the pontine nuclei, mPFC neurons that project various cortical and subcortical areas^82, 83, 84^ have substantial roles in facilitating rapid adaptation. A particularly noteworthy structure involved in this process is the thalamo-cortical pathway. This ability of the thalamo-cortical circuit to signal rule switching has been observed in other tasks that demand flexible adjustments in both cue-specific and task-specific activities^28, 85, 86, 87, 88^. Interestingly, a subset of mPFC neurons showed a sustained increase in their baseline firing rates in response to the paradigm adaptations (Fig. 8). This sustained increase may contribute to the heightened gamma band (30-80 Hz) oscillation in the mPFC during behavior, a phenomenon associated with enhanced attention and cognitive flexibility^78, 82, 89^. Moreover, the mPFC might regulate CR onset timing by balancing motor activation and inhibition^90, 91, 92^. During DEC-to-TEC adaptation, animals are required to withhold action, while during TEC-to-DEC adaptation, they need to actively reduce the action delay. The dynamic equilibrium between proactive and reactive action planning, orchestrated by mPFC signaling, provides the flexibility for controlling action timing. It remains unclear which downstream pathways of the thalamo-cortical loop are involved in the rapid temporal adaptation of EBC. Future research should delve into exploring the neuronal mechanisms of communication across multiple brain regions and their collective roles in regulating adaptable sensorimotor behaviors.

## Methods

### Animals

Wild type C57Bl/6 mice aged from 12 to 20 weeks were included in the experiments (both male and female) and housed individually (12:12 light/dark cycle) with food and water *ad libitum*. The experiments were approved by the institutional animal welfare committee of Erasmus MC. In total, 24 mice were trained with the TEC paradigm and 20 mice were trained with the DEC paradigm for electrophysiological recording and behavior manipulation. We also included a dataset of IpN recordings in seven DEC-trained mice from our previous study^11^.

### Surgery

Mice were anesthetized with 5% isoflurane for induction, 2% isoflurane for maintenance, and kept at 37°C using a heating pad during surgery. After fixing the animal into a standard mouse stereotaxic surgical plate (Stoelting Co., Wood Dale IL, USA), we exposed the skull and applied Optibond All-In-One (Kerr, Scafati SA, Italy) for better pedestal attachment. A small brass pedestal was attached onto the skull and fixed with Charisma (Heraeus Kulzer GmbH, Hanau, Germany). We performed craniotomy (approximately 1.5 mm diameter) above the target region (IpN or mPFC) for electrophysiology, electrical stimulation, and muscimol injection. The stereotaxic coordinates for the craniotomy over the IpN are: AP: −2.2 mm (measured from Lambda), ML: 1.8 mm; the craniotomy over the mPFC: AP: +1.9 mm (measured from Bregma), ML: 0.4 mm. A small chamber was made by attaching Charisma around the skull of craniotomy, and finally closed by a low-viscosity elastomer sealant (Kwik-cast, World Precision Instruments, Sarasota FL, USA). Bupivacaine hydrochloride (2.5 mg mL^−1^, *i.p.*) was administrated after surgery.

### Behavioral training

After one-week recovery from the surgery, mice were head-fixed on a cylindrical treadmill in a sound- and light-attenuated setup for one-week habituation^7^. For DEC and TEC training, a green LED (CS) was placed approximately 7 cm in front of the mouse, while a corneal air-puff (US, tip opening Φ = 0.8 mm, 30 psi) was directed at the left eye with 5 mm distance. The CS duration was 250 ms in both TEC and DEC paradigms. A 15-ms US air-puff was delivered at the CS offset for the DEC paradigm but with 250 ms interval after CS offset for the TEC paradigm. Randomized inter trial intervals of 8-12 seconds were employed, with a minimum of 200 trials per training session. We introduced CS-only trials randomly within the normal paired trials at a 1:6 ratio. Animals achieving a high CR-trial probability (70% of total trials) were considered as well-trained. Eyelid movement was captured by a 250-fps camera (scA640-120gm, Basler, Ahrensburg, Germany). National Instruments System (NI 9263 and NI 9269, National Instruments, Austin TX, USA) was used to control the triggers and digitization of camera signal using custom Labview codes. The RHD2000 Evaluation System (Intan Technology) with a 20 kHz sample rate recorded digitized triggers and eyelid positions.

### In-vivo electrophysiology

After a recovery period of 2-3 days from the craniotomy, we conducted *in-vivo* electrophysiological recordings on awake behaving mice. We performed acute recordings throughout the study using acute 64-channel silicon probes (ASSY 77H2, Cambridge NeuroTech). We vertically inserted it into either the IpN (at approximately 2.4 mm depth) or mPFC (at approximately 1.5 mm depth), guided by an electrode manipulator (Luigs and Neumann SM7, Germany). Neuronal signals were notch-filtered at 50 Hz, amplified, and digitized by an Intan RHD2000 Evaluation System (Intan Technology) at 20 kHz sampling rate and were further analyzed offline using custom-written Matlab codes. For the electrophysiological experiments during TEC-to-DEC and DEC-to-TEC adaptation, we recorded the activities of same neurons for at least 50 trials prior to and after paradigm switch. In general, we recorded 5 days for each recording region for all types of behavior paradigms, except for the adaptation paradigms, which was 2 days to avoid long term learning.

### IpN electrostimulation

Data was acquired according to our previous study^12^. Briefly, after IpN craniotomy, a stimulation glass electrode (8 μm tip opening) with 2 M saline-0.5% alcian blue was penetrated into IpN. Electrical stimuli (500 Hz with 250 μs biphasic pulses and 200 ms pulse train) were performed to the animals with current strength of 0.6, 0.8, 1.0, and 1.2 μA. Eyelid movement ipsilateral to the stimulation side was recorded using the devices mentioned above. The maximum eyelid closure during 1.2 μA stimulation was viewed as full eyelid closure.

### Pharmacological inhibition of mPFC

For mPFC inhibition during TEC-to-DEC adaptation, a small craniotomy was made over bilateral mPFC in TEC trained mouse. We first recorded 50 TEC trials as control, then gently injected 20 nL muscimol (0.5 mg/mL, Tocris Bioscience, 0289) in the mPFC. Five minutes later, we switched to DEC paradigm to test their CR adaptation. We performed the similar procedure for the DEC-to-TEC adaptation experiment.

### Histology and imaging

Animals were deeply anesthetized by injecting pentobarbital sodium solution (50 mg/kg, *i.p.*) and transcardially perfused with saline and the solution of 4% paraformaldehyde (PFA) in 0.1 M phosphate buffer (PB, pH = 0.74). The brains were removed immediately and post-fixed in 4% PFA for 2 hours, and then dehydrated by 10% sucrose in 4°C fridge overnight. On the next day, the brains were embedded in 12% gelatin with 10% sucrose, fixed again in 10% formalin-30% sucrose solution for two hours, and then dehydrated with 30% sucrose 4°C overnight. Coronal brain slices with a thickness of 50 μm were obtained using a microtome (SM2000R, Leica), and collected in 0.1 M PB. Brain slices with DiI labeling were stained with DAPI as background and imaged by Zeiss LSM700. Slices with alcian blue labeling were stained with neutral red as background and scanned by NanoZoomer (Hamamatsu).

### Eyeblink data analysis

As described in previous work^11, 12^, the investigation of eyelid position changes in response to the CS and US involved normalizing each trial to a 500 ms baseline preceding the CS onset. We removed trials with noisy baseline (spontaneous blinking) by performing an iterative Grubbs’ outlier detection test (α = 0.05) on the standard deviations of baseline. A CR trial was determined if the eyelid closure exceeded 5% of the mean baseline, and CR onset was defined as the timing with which eyelid closure exceeded six standard deviations of the baseline value. We also explored alternative criteria for detecting CR onset, based on CR amplitude^11^ and CR velocity^39^ from prior studies. Peak amplitudes of CR and UR were detected as the maximum during the CS-US interval and a 100 ms window after US, respectively. CR slope was defined as the angle of a linear fit between 10% and 90% of CR peak amplitudes. Maximum CR velocity was calculated with 10 ms bins.

### Electrophysiology analysis

We analyzed the electrophysiological recordings using custom Matlab code. Raw recordings were band-pass filtered between 300-3000 Hz to subtract noise and field potential signals. We extracted spike events with amplitudes that exceed three SDs of the baseline value. Multichannel recordings were sorted using JRCLUST^89, 90^, and all spike times were stored for further analysis. We transformed each spike into a Gaussian kernel (21 ms kernel for IpN single units, and 41 ms kernel for mPFC single units) and aligned with CS onset for the peristimulus time histograms (PSTH) calculation. Single trial instantaneous firing rate was further smoothed by another 10 ms sliding window for trial-by-trial analysis. To ensure data reliability, only cells with over 20 CR trials were included in the dataset for analysis of cell modulation in response to the CS. For IpN neurons, CR-related modulation was detected from 10 ms after CS onset to 10 ms before US onset, considering potential epoch crossings in PSTH construction. Baseline firing rate was calculated as the mean frequency in a 500 ms window prior to the CS onset. Neurons with PSTH changes exceeding three SDs of the baseline frequency were considered as CR-related modulation neurons, and the first time point reaching to the 3-SD threshold was the modulation onset. For mPFC neurons, we determined CR modulation by checking the significance of the baseline firing rate to the firing rate between CS and US. Transient mPFC facilitation neurons were characterized as cells with a half-wave width constituting less than 20% of the CS-US interval, along with a modulation peak occurring prior to 200 ms. The remaining modulated mPFC neurons were classified as part of the sustained facilitation group.

### Decoding classifier analysis

The decoding classifier analysis was adapted from the neural decoding toolbox, as described previously^41^. We utilized the max correlation coefficient classifier to decode potential differences in firing patterns of IpN single units between early and late CR onset trials in TEC recordings. To execute this, we organized all trials from a single recording session based on their CR onset times, evenly splitting them into early- and late-onset subgroups. Within a time window ranging from −500 to 500 ms, we established bins of 100 ms with a step of 50 ms to cluster spikes. In each decoding cycle, 90% of the trials were randomly selected from both early and late groups as training datasets, and the rest 10% of both became as testing dataset. This process was repeated 20 times as resampling. The accuracy of decoding was calculated by determining the percentage of accurately distinguishing early and late trials within the testing dataset. To assess the statistical significance of the decoding outcome, this procedure was repeated 10 times using shuffled data. IpN neurons that exhibited a decoding difference of at least 100 ms in modulation peak times, while maintaining less than a 20% difference in modulation amplitude, were categorized as temporal modulation neurons (Extended Data Fig. 4b). Neurons with amplitude differences exceeding 20% were classified as amplitude modulation neurons (Extended Data Fig. 4c, d).

### Statistics

All the analysis and statistics were performed using MATLAB and GraphPad Prism. The average eyelid closure curve in was plotted as mean ± standard deviation (SD), other data were demonstrated as mean ± standard error of mean (s.e.m.). Wilcoxon signed-rank test was used to detect the group differences of paired samples, except for the IpN modulation onset comparison in DEC-to-TEC and TEC-to-DEC adaptation experiments, which paired *t*-test was applied. Mann-Whitney U test was performed to compare the group differences of independent samples. Repeated measures one-way ANOVA was used for analyzing the data continuously collected from the same subject. Pearson’s r was used to detect the correlation between two continuously distributed variables. All the statistical tests were two-tail based. *P* ≤ 0.05 was considered as significantly different, and were annotated as *: *P* ≤ 0.05, **: *P* ≤ 0.01, ***: *P* ≤ 0.001, and ****: *P* ≤ 0.0001.

## Acknowledgement

The authors thank M. Rutteman (Department of Neuroscience, Erasmus MC) for managing the animals, S. Dijkhuizen (Department of Neuroscience, Erasmus MC) for providing the eyeblink conditioning training boxes, S. Yu (Department of Neuroscience, Erasmus MC) for technical support of the electrophysiology setup and MATLAB programming, H. Hasanbegovic, and C. Schafer (Department of Neuroscience, Erasmus MC) for the constructive inputs to this study. This work is supported by CSC fellowship (Z. R., CSC201907720092), NWO VIDI grant (Z.G., VI.Vidi.192.008), NWO-Klein grant (Z.G., OCENW.KLEIN.007) and ERC-stg grant (Z.G., 852869).

## Author contributions

Z.R. and Z.G. conceived the project. Z.R. performed most of the experiments and analysis. M.A. contributed to mice training and electrophysiology recording in the study. X.W. provided IpN electrophysiology data in DEC trained mice and IpN microsimulation data. Z.R., C.I.D.Z., and Z.G. wrote the whole manuscript with inputs from all the authors. Z.G. supervised the whole project.

## References

1. Rescorla RA. Probability of shock in the presence and absence of CS in fear conditioning. Journal of comparative and physiological psychology 66, 1–5 (1968).

2. Johansen JP, Cain CK, Ostroff LE, LeDoux JE. Molecular mechanisms of fear learning and memory. Cell 147, 509–524 (2011).

3. Guo ZV, et al. Flow of cortical activity underlying a tactile decision in mice. Neuron 81, 179–194 (2014).

4. Li N, Chen TW, Guo ZV, Gerfen CR, Svoboda K. A motor cortex circuit for motor planning and movement. Nature 519, 51–56 (2015).

5. Goard MJ, Pho GN, Woodson J, Sur M. Distinct roles of visual, parietal, and frontal motor cortices in memory-guided sensorimotor decisions. eLife 5, (2016).

6. Halverson HE, Khilkevich A, Mauk MD. Cerebellar Processing Common to Delay and Trace Eyelid Conditioning. The Journal of neuroscience : the official journal of the Society for Neuroscience 38, 7221–7236 (2018).

7. Chettih SN, McDougle SD, Ruffolo LI, Medina JF. Adaptive timing of motor output in the mouse: the role of movement oscillations in eyelid conditioning. Front Integr Neurosci 5, 72 (2011).

8. Siegel JJ, et al. Trace Eyeblink Conditioning in Mice Is Dependent upon the Dorsal Medial Prefrontal Cortex, Cerebellum, and Amygdala: Behavioral Characterization and Functional Circuitry. eNeuro 2, (2015).

9. Heiney SA, Wohl MP, Chettih SN, Ruffolo LI, Medina JF. Cerebellar-dependent expression of motor learning during eyeblink conditioning in head-fixed mice. The Journal of neuroscience : the official journal of the Society for Neuroscience 34, 14845–14853 (2014).

10. ten Brinke MM, et al. Evolving Models of Pavlovian Conditioning: Cerebellar Cortical Dynamics in Awake Behaving Mice. Cell reports 13, 1977–1988 (2015).

11. Ten Brinke MM, et al. Dynamic modulation of activity in cerebellar nuclei neurons during pavlovian eyeblink conditioning in mice. eLife 6, (2017).

12. Wang X, Yu SY, Ren Z, De Zeeuw CI, Gao Z. A FN-MdV pathway and its role in cerebellar multimodular control of sensorimotor behavior. Nature communications 11, 6050 (2020).

13. Delgado-Garcia JM, Gruart A. Firing activities of identified posterior interpositus nucleus neurons during associative learning in behaving cats. Brain Res Brain Res Rev 49, 367–376 (2005).

14. Takehara-Nishiuchi K. The Anatomy and Physiology of Eyeblink Classical Conditioning. Curr Top Behav Neurosci 37, 297–323 (2018).

15. Freeman JH, Steinmetz AB. Neural circuitry and plasticity mechanisms underlying delay eyeblink conditioning. Learn Mem 18, 666–677 (2011).

16. Yang Y, Lei C, Feng H, Sui JF. The neural circuitry and molecular mechanisms underlying delay and trace eyeblink conditioning in mice. Behav Brain Res 278, 307–314 (2015).

17. McCormick DA, Thompson RF. Neuronal responses of the rabbit cerebellum during acquisition and performance of a classically conditioned nictitating membrane-eyelid response. The Journal of neuroscience : the official journal of the Society for Neuroscience 4, 2811–2822 (1984).

18. Gonzalez-Joekes J, Schreurs BG. Anatomical characterization of a rabbit cerebellar eyeblink premotor pathway using pseudorabies and identification of a local modulatory network in anterior interpositus. The Journal of neuroscience : the official journal of the Society for Neuroscience 32, 12472–12487 (2012).

19. Flumerfelt BA, Otabe S, Courville J. Distinct projections to the red nucleus from the dentate and interposed nuclei in the monkey. Brain Res 50, 408–414 (1973).

20. Teune TM, van der Burg J, van der Moer J, Voogd J, Ruigrok TJ. Topography of cerebellar nuclear projections to the brain stem in the rat. Prog Brain Res 124, 141–172 (2000).

21. Aksenov D, Serdyukova N, Irwin K, Bracha V. GABA neurotransmission in the cerebellar interposed nuclei: involvement in classically conditioned eyeblinks and neuronal activity. Journal of neurophysiology 91, 719–727 (2004).

22. Parker KL, Zbarska S, Carrel AJ, Bracha V. Blocking GABAA neurotransmission in the interposed nuclei: effects on conditioned and unconditioned eyeblinks. Brain Res 1292, 25–37 (2009).

23. Jirenhed DA, Rasmussen A, Johansson F, Hesslow G. Learned response sequences in cerebellar Purkinje cells. Proc Natl Acad Sci U S A 114, 6127–6132 (2017).

24. Gruart A, Guillazo-Blanch G, Fernandez-Mas R, Jimenez-Diaz L, Delgado-Garcia JM. Cerebellar posterior interpositus nucleus as an enhancer of classically conditioned eyelid responses in alert cats. Journal of neurophysiology 84, 2680–2690 (2000).

25. McCormick DA, Clark GA, Lavond DG, Thompson RF. Initial localization of the memory trace for a basic form of learning. Proc Natl Acad Sci U S A 79, 2731–2735 (1982).

26. Jimenez-Diaz L, Navarro-Lopez Jde D, Gruart A, Delgado-Garcia JM. Role of cerebellar interpositus nucleus in the genesis and control of reflex and conditioned eyelid responses. The Journal of neuroscience : the official journal of the Society for Neuroscience 24, 9138–9145 (2004).

27. Isoda M, Hikosaka O. Switching from automatic to controlled action by monkey medial frontal cortex. Nature neuroscience 10, 240–248 (2007).

28. Wang J, Narain D, Hosseini EA, Jazayeri M. Flexible timing by temporal scaling of cortical responses. Nature neuroscience 21, 102–110 (2018).

29. Kronforst-Collins MA, Disterhoft JF. Lesions of the caudal area of rabbit medial prefrontal cortex impair trace eyeblink conditioning. Neurobiol Learn Mem 69, 147–162 (1998).

30. Weible AP, McEchron MD, Disterhoft JF. Cortical involvement in acquisition and extinction of trace eyeblink conditioning. Behavioral neuroscience 114, 1058–1067 (2000).

31. McLaughlin J, Skaggs H, Churchwell J, Powell DA. Medial prefrontal cortex and pavlovian conditioning: trace versus delay conditioning. Behavioral neuroscience 116, 37–47 (2002).

32. Takehara-Nishiuchi K, Kawahara S, Kirino Y. NMDA receptor-dependent processes in the medial prefrontal cortex are important for acquisition and the early stage of consolidation during trace, but not delay eyeblink conditioning. Learn Mem 12, 606–614 (2005).

33. Siegel JJ, Kalmbach B, Chitwood RA, Mauk MD. Persistent activity in a cortical-to-subcortical circuit: bridging the temporal gap in trace eyelid conditioning. Journal of neurophysiology 107, 50–64 (2012).

34. Weible AP, Weiss C, Disterhoft JF. Activity profiles of single neurons in caudal anterior cingulate cortex during trace eyeblink conditioning in the rabbit. Journal of neurophysiology 90, 599–612 (2003).

35. Siegel JJ, Mauk MD. Persistent activity in prefrontal cortex during trace eyelid conditioning: dissociating responses that reflect cerebellar output from those that do not. The Journal of neuroscience : the official journal of the Society for Neuroscience 33, 15272–15284 (2013).

36. Hattori S, Yoon T, Disterhoft JF, Weiss C. Functional reorganization of a prefrontal cortical network mediating consolidation of trace eyeblink conditioning. The Journal of neuroscience : the official journal of the Society for Neuroscience 34, 1432–1445 (2014).

37. Wu GY, et al. Optogenetic Inhibition of Medial Prefrontal Cortex-Pontine Nuclei Projections During the Stimulus-free Trace Interval Impairs Temporal Associative Motor Learning. Cerebral cortex 28, 3753–3763 (2018).

38. Chen H, et al. Prefrontal control of cerebellum-dependent associative motor learning. Cerebellum 13, 64–78 (2014).

39. Ohmae S, Medina JF. Climbing fibers encode a temporal-difference prediction error during cerebellar learning in mice. Nature neuroscience 18, 1798–1803 (2015).

40. Ohyama T, Nores WL, Medina JF, Riusech FA, Mauk MD. Learning-induced plasticity in deep cerebellar nucleus. The Journal of neuroscience : the official journal of the Society for Neuroscience 26, 12656–12663 (2006).

41. Meyers EM. The neural decoding toolbox. Front Neuroinform 7, 8 (2013).

42. Plakke B, Freeman JH, Poremba A. Metabolic mapping of the rat cerebellum during delay and trace eyeblink conditioning. Neurobiol Learn Mem 88, 11–18 (2007).

43. Green JT, Arenos JD. Hippocampal and cerebellar single-unit activity during delay and trace eyeblink conditioning in the rat. Neurobiol Learn Mem 87, 269–284 (2007).

44. Berthier NE, Moore JW. Activity of deep cerebellar nuclear cells during classical conditioning of nictitating membrane extension in rabbits. Exp Brain Res 83, 44–54 (1990).

45. Kalmbach BE, Ohyama T, Kreider JC, Riusech F, Mauk MD. Interactions between prefrontal cortex and cerebellum revealed by trace eyelid conditioning. Learning & memory 16, 86–95 (2009).

46. Narain D, Remington ED, Zeeuw CI, Jazayeri M. A cerebellar mechanism for learning prior distributions of time intervals. Nature communications 9, 469 (2018).

47. MacDonald CJ, Lepage KQ, Eden UT, Eichenbaum H. Hippocampal “time cells” bridge the gap in memory for discontiguous events. Neuron 71, 737–749 (2011).

48. Tsutsumi S, et al. Purkinje Cell Activity Determines the Timing of Sensory-Evoked Motor Initiation. Cell reports 33, 108537 (2020).

49. Ito M. Cerebellar long-term depression: characterization, signal transduction, and functional roles. Physiological reviews 81, 1143–1195 (2001).

50. Ohmae S, Ohmae K, Heiney S, Subramanian D, Medina J. A recurrent circuit links antagonistic cerebellar modules during associative motor learning. bioRxiv, 2021.2011.2016.468438 (2021).

51. Weiss C, Disterhoft JF. Exploring prefrontal cortical memory mechanisms with eyeblink conditioning. Behavioral neuroscience 125, 318–326 (2011).

52. Morcuende S, Delgado-Garcia JM, Ugolini G. Neuronal premotor networks involved in eyelid responses: retrograde transneuronal tracing with rabies virus from the orbicularis oculi muscle in the rat. The Journal of neuroscience : the official journal of the Society for Neuroscience 22, 8808–8818 (2002).

53. Kim JJ, Clark RE, Thompson RF. Hippocampectomy impairs the memory of recently, but not remotely, acquired trace eyeblink conditioned responses. Behavioral neuroscience 109, 195–203 (1995).

54. Woodruff-Pak DS, Disterhoft JF. Where is the trace in trace conditioning? Trends in neurosciences 31, 105–112 (2008).

55. Giovannucci A, et al. Cerebellar granule cells acquire a widespread predictive feedback signal during motor learning. Nature neuroscience 20, 727–734 (2017).

56. Gao Z, et al. Excitatory Cerebellar Nucleocortical Circuit Provides Internal Amplification during Associative Conditioning. Neuron 89, 645–657 (2016).

57. Houck BD, Person AL. Cerebellar Premotor Output Neurons Collateralize to Innervate the Cerebellar Cortex. The Journal of comparative neurology 523, 2254–2271 (2015).

58. Lanore F, Cayco-Gajic NA, Gurnani H, Coyle D, Silver RA. Cerebellar granule cell axons support high-dimensional representations. Nature neuroscience 24, 1142–1150 (2021).

59. Boele HJ, Koekkoek SK, De Zeeuw CI, Ruigrok TJ. Axonal sprouting and formation of terminals in the adult cerebellum during associative motor learning. The Journal of neuroscience : the official journal of the Society for Neuroscience 33, 17897–17907 (2013).

60. Pugh JR, Raman IM. Potentiation of mossy fiber EPSCs in the cerebellar nuclei by NMDA receptor activation followed by postinhibitory rebound current. Neuron 51, 113–123 (2006).

61. Mauk MD, Ruiz BP. Learning-dependent timing of Pavlovian eyelid responses: differential conditioning using multiple interstimulus intervals. Behavioral neuroscience 106, 666–681 (1992).

62. Kalmbach BE, Ohyama T, Mauk MD. Temporal patterns of inputs to cerebellum necessary and sufficient for trace eyelid conditioning. Journal of neurophysiology 104, 627–640 (2010).

63. Gao Z, van Beugen BJ, De Zeeuw CI. Distributed synergistic plasticity and cerebellar learning. Nature reviews Neuroscience 13, 619–635 (2012).

64. Allen WE, et al. Global Representations of Goal-Directed Behavior in Distinct Cell Types of Mouse Neocortex. Neuron 94, 891–907 e896 (2017).

65. Leal-Campanario R, Delgado-Garcia JM, Gruart A. The rostral medial prefrontal cortex regulates the expression of conditioned eyelid responses in behaving rabbits. The Journal of neuroscience : the official journal of the Society for Neuroscience 33, 4378–4386 (2013).

66. Wu GY, et al. Reevaluating the role of the medial prefrontal cortex in delay eyeblink conditioning. Neurobiology of learning and memory 97, 277–288 (2012).

67. Wu GY, et al. Medial Prefrontal Cortex-Pontine Nuclei Projections Modulate Suboptimal Cue-Induced Associative Motor Learning. Cerebral cortex 28, 880–893 (2018).

68. Arnsten AF. Stress weakens prefrontal networks: molecular insults to higher cognition. Nature neuroscience 18, 1376–1385 (2015).

69. Arnsten AF. Stress signalling pathways that impair prefrontal cortex structure and function. Nature reviews Neuroscience 10, 410–422 (2009).

70. Giustino TF, Maren S. Noradrenergic Modulation of Fear Conditioning and Extinction. Frontiers in behavioral neuroscience 12, 43 (2018).

71. Albergaria C, Silva NT, Pritchett DL, Carey MR. Locomotor activity modulates associative learning in mouse cerebellum. Nature neuroscience 21, 725–735 (2018).

72. Shors TJ, Weiss C, Thompson RF. Stress-induced facilitation of classical conditioning. Science 257, 537–539 (1992).

73. Bolkan SS, et al. Thalamic projections sustain prefrontal activity during working memory maintenance. Nature neuroscience 20, 987–996 (2017).

74. Abbas AI, et al. Somatostatin Interneurons Facilitate Hippocampal-Prefrontal Synchrony and Prefrontal Spatial Encoding. Neuron 100, 926–939 e923 (2018).

75. Kim D, et al. Distinct Roles of Parvalbumin- and Somatostatin-Expressing Interneurons in Working Memory. Neuron 92, 902–915 (2016).

76. Liu D, et al. Medial prefrontal activity during delay period contributes to learning of a working memory task. Science 346, 458–463 (2014).

77. Spellman T, Rigotti M, Ahmari SE, Fusi S, Gogos JA, Gordon JA. Hippocampal-prefrontal input supports spatial encoding in working memory. Nature 522, 309–314 (2015).

78. Kim H, Ahrlund-Richter S, Wang X, Deisseroth K, Carlen M. Prefrontal Parvalbumin Neurons in Control of Attention. Cell 164, 208–218 (2016).

79. White MG, et al. Anterior Cingulate Cortex Input to the Claustrum Is Required for Top-Down Action Control. Cell reports 22, 84–95 (2018).

80. Le Merre P, et al. Reward-Based Learning Drives Rapid Sensory Signals in Medial Prefrontal Cortex and Dorsal Hippocampus Necessary for Goal-Directed Behavior. Neuron 97, 83–91 e85 (2018).

81. Pinto L, Dan Y. Cell-Type-Specific Activity in Prefrontal Cortex during Goal-Directed Behavior. Neuron 87, 437–450 (2015).

82. Le Merre P, Ahrlund-Richter S, Carlen M. The mouse prefrontal cortex: Unity in diversity. Neuron 109, 1925–1944 (2021).

83. Harris JA, et al. Hierarchical organization of cortical and thalamic connectivity. Nature 575, 195–202 (2019).

84. Gao L, et al. Single-neuron projectome of mouse prefrontal cortex. Nature neuroscience 25, 515–529 (2022).

85. Banerjee A, Parente G, Teutsch J, Lewis C, Voigt FF, Helmchen F. Value-guided remapping of sensory cortex by lateral orbitofrontal cortex. Nature 585, 245–250 (2020).

86. Rikhye RV, Gilra A, Halassa MM. Thalamic regulation of switching between cortical representations enables cognitive flexibility. Nature neuroscience 21, 1753–1763 (2018).

87. Spellman T, Svei M, Kaminsky J, Manzano-Nieves G, Liston C. Prefrontal deep projection neurons enable cognitive flexibility via persistent feedback monitoring. Cell 184, 2750–2766 e2717 (2021).

88. de Kloet SF, et al. Bi-directional regulation of cognitive control by distinct prefrontal cortical output neurons to thalamus and striatum. Nature communications 12, 1994 (2021).

89. Cho KKA, Davidson TJ, Bouvier G, Marshall JD, Schnitzer MJ, Sohal VS. Cross-hemispheric gamma synchrony between prefrontal parvalbumin interneurons supports behavioral adaptation during rule shift learning. Nature neuroscience 23, 892–902 (2020).

90. Eagle DM, Baunez C, Hutcheson DM, Lehmann O, Shah AP, Robbins TW. Stop-signal reaction-time task performance: role of prefrontal cortex and subthalamic nucleus. Cerebral cortex 18, 178–188 (2008).

91. Baunez C, Salin P, Nieoullon A, Amalric M. Impaired performance in a conditioned reaction time task after thermocoagulatory lesions of the fronto-parietal cortex in rats. Cerebral cortex 8, 301–309 (1998).

92. Hardung S, et al. A Functional Gradient in the Rodent Prefrontal Cortex Supports Behavioral Inhibition. Current biology : CB 27, 549–555 (2017).

