## Extended data Figures 1-12 for "Neuronal dynamics of cerebellum and medial prefrontal cortex in adaptive motor timing"

**Ren *et al.***

**Extended data Figures 1-12**

### Extended data Figure 1

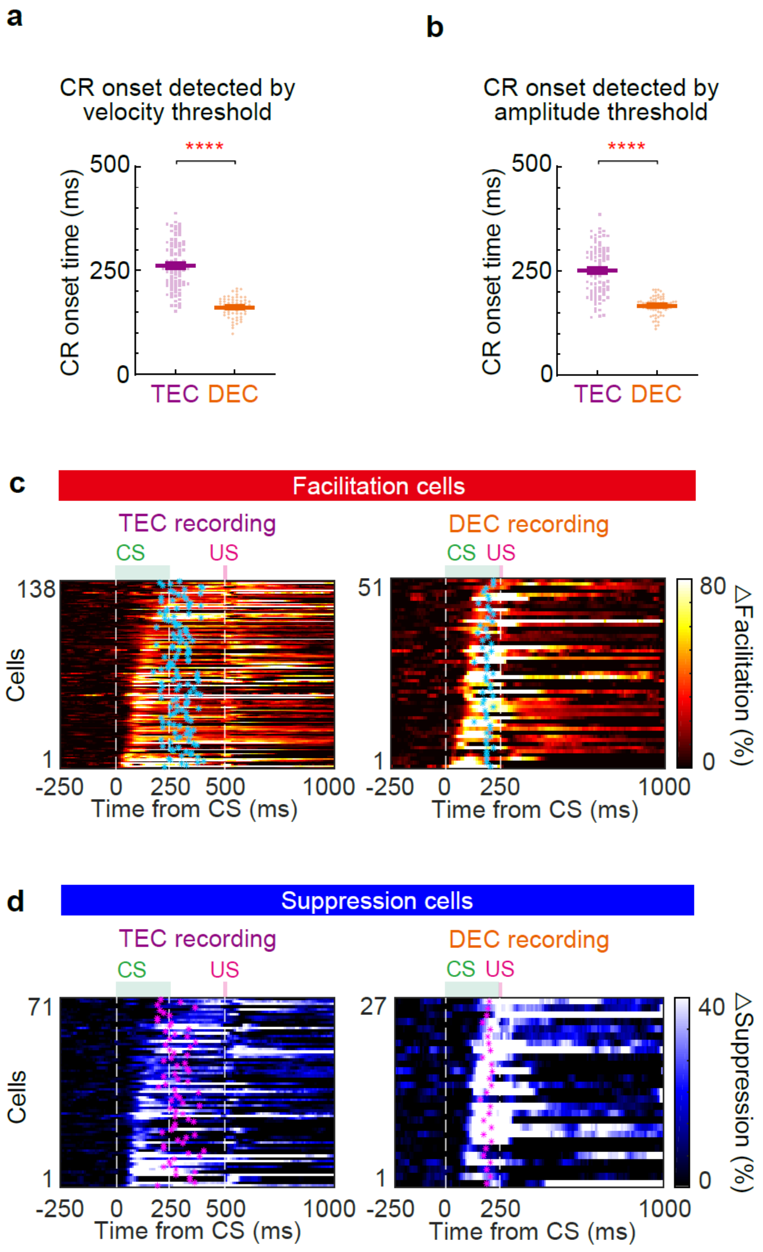


**Extended data Figure 1: Summary of IpN neuron recordings during TEC and DEC. Data related to fig.1.**
**a, b** The CR onset time of TEC and DEC recordings which was detected by CR velocity (**a**) and CR amplitude threshold (**b**) ($P$ < 0.0001 for both, $n$ = 92 and 67 sessions). **c** Summary of IpN neurons that had increased spike rates (facilitation cells) during TEC ($n$ = 138 cells, left) and DEC ($n$ = 51 cells, right). Each row of the heatmap represents normalized spike rate of one IpN neuron. Blue dots represent the mean CR onset timing of the corresponding session. **d** Same as (**c**), but for the IpN neurons that had decreased spike rates (suppression cells, $n$ = 71 and 27 cells for TEC and DEC

respectively). Magenta dots represent the mean CR onsets of the corresponding sessions.

### Extended data Figure 2

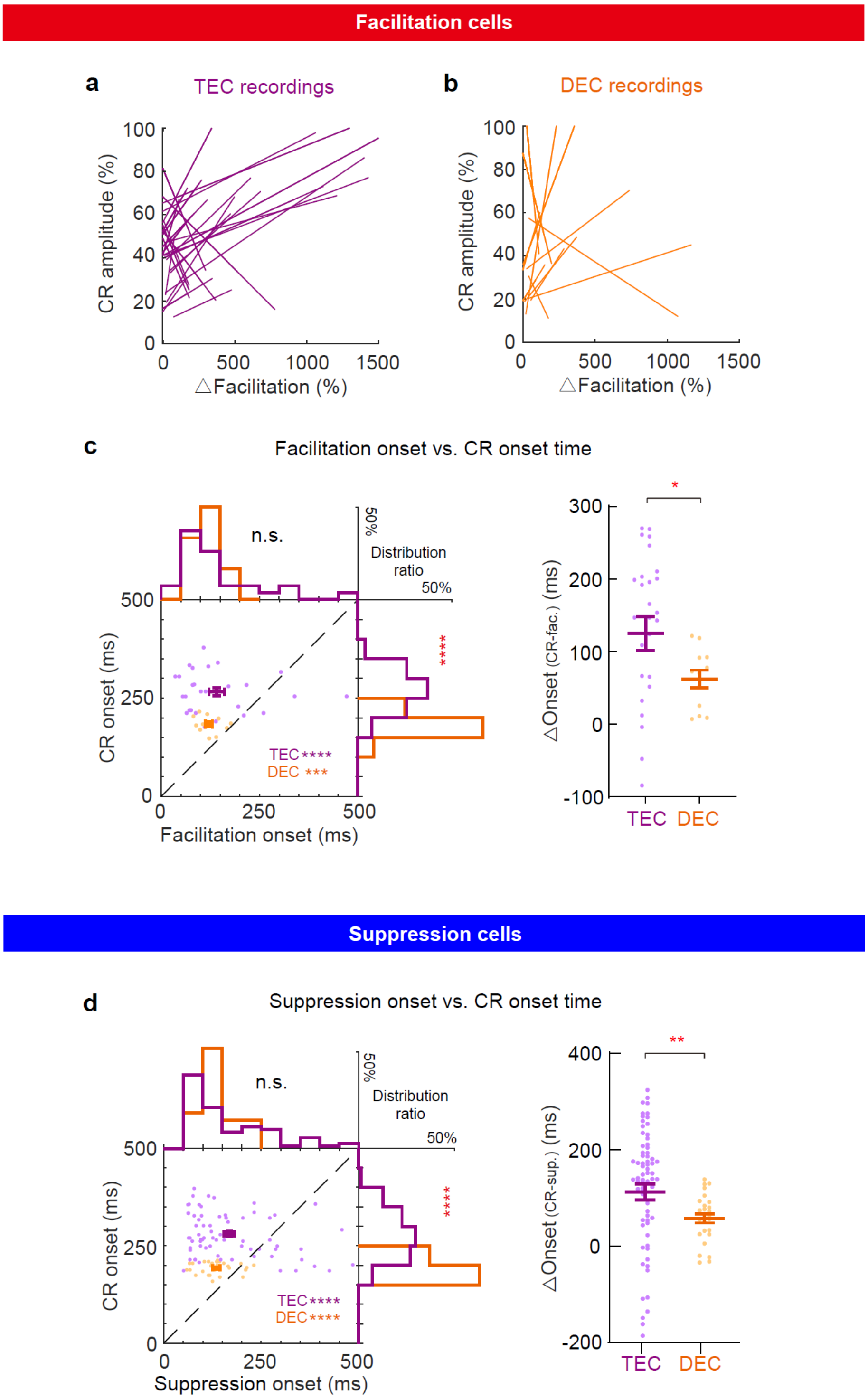


**Extended data Figure 2: Temporal dynamics of IpN neuron activity and CRs in response to different CS-US intervals. Data related to fig.2.**

**a, b** Trial-by-trial correlation between IpN neuron activity and CR amplitude in TEC trained mice (*n* = 27 cells) and DEC trained mice (*n* = 12 cells). Each line represents the linear fit of the trial-by-trial correlation from one neuron ($P$ < 0.05). See Methods for detailed calculation. **c** Summary of the facilitation onset timing and the corresponding CR onset timing during TEC and DEC (recordings are from **a, b**). Left: scatter plots and histograms illustrating the relationship of onset timings IpN facilitation and CRs. IpN neuron modulation precedes CR onset in both TEC and DEC ($P = 6.33 \times 10^{-5}$ for TEC, $P = 0.00049$ for DEC; comparison of CR onsets of TEC and DEC: $P = 5.60 \times 10^{-7}$; facilitation onsets during TEC and DEC: $P = 0.51$, $n = 27$ and cells). Right: time interval between the onset of IpN activity and the onset of CR (△onset, CR onsets minus IpN neuron modulation onsets) during TEC and DEC, $P = 0.026$, $n = 27$ and 12 cells. **d** Same as (**c**), but for suppression IpN cells. Comparison of the CR onsets and IpN suppression onsets. $P = 7.32 \times 10^{-9}$ for TEC, $P = 4.57 \times 10^{-6}$ for DEC; comparison of IpN suppression onsets during TEC and DEC, $P = 0.68$; CR onsets of TEC and DEC, $P = 1.0 \times 10^{-15}$; TEC and DEC △onset: $P = 0.0010$, $n = 71$ and 27 cells. Data is shown as mean ± s.e.m., n.s., not significant, *$P \leq 0.05$, **$P \leq 0.01$, ***$P \leq 0.001$, and ****$P \leq 0.0001$.

### Extended data Figure 3

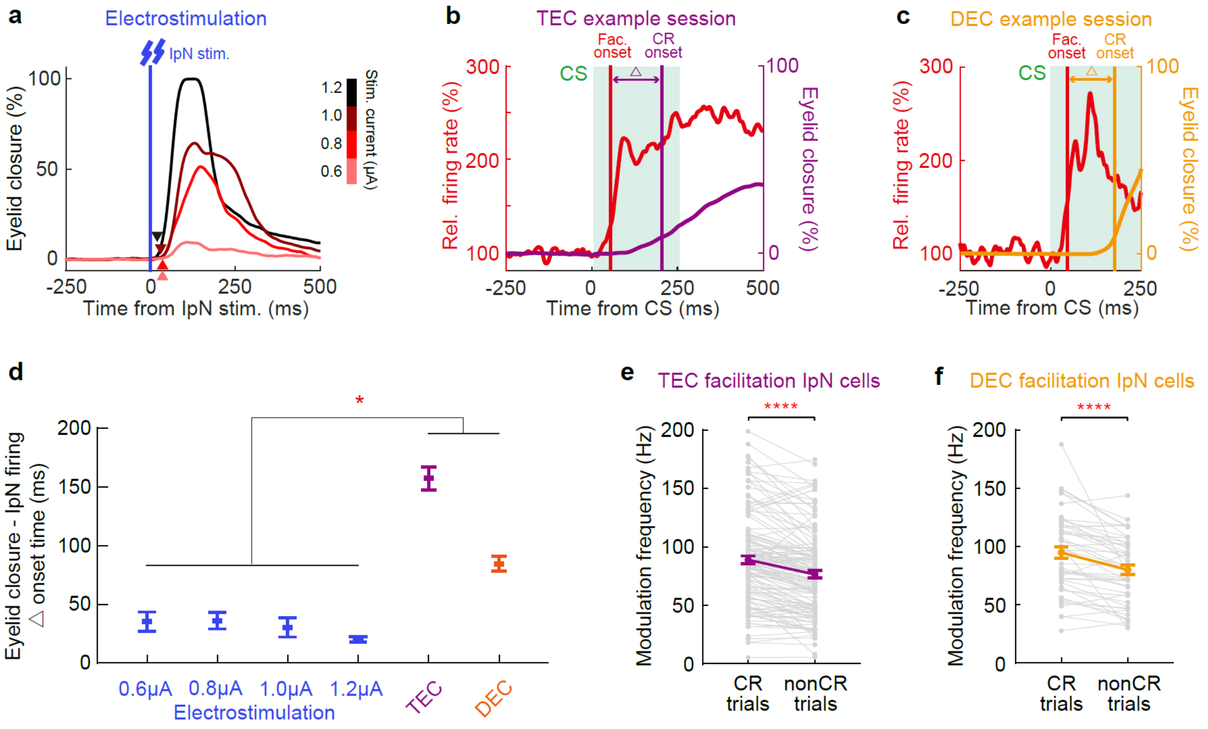


**Extended data Figure 3: Eyelid closure induced by electrostimulation of IpN has shorter latency comparing to associative eyelid closure in TEC and DEC. Data related to fig.2.**
**a** Eyelid traces in response to IpN electrostimulations. Stimulation intensities are indicated on the right. Arrowheads indicate the onsets of eyelid closure. **b** Spike rate (red) and eyelid closure (purple) from an example IpN neuron recording during TEC. Vertical lines indicate the facilitation onset and CR onset. **c** Same as (**b**), but from an example IpN neuron recorded during DEC. **d** delays between IpN modulation (Δonset) and CR are much longer than the delay between electrostimulation and eyelid closure (comparison of Δonset between electrostimulations and TEC/DEC, *P* = 0.024, 0.026, 0.021, and 0.013 for TEC versus electric stimulations (four intensities); *P* = 0.041, 0.046, 0.033, and 0.019 for DEC versus four electric stimulation). **e-f** Comparison of CS-related modulation frequency between CR and non-CR trials in all facilitation IpN cells recorded in TEC- (**e**) or DEC- (**f**) trained mice (*P* < 0.0001 for both, *n* = 138 and 48 cells). Data is shown as the mean ± s.e.m., **P* ≤ 0.05, and *****P* ≤ 0.0001.

### Extended data Figure 4

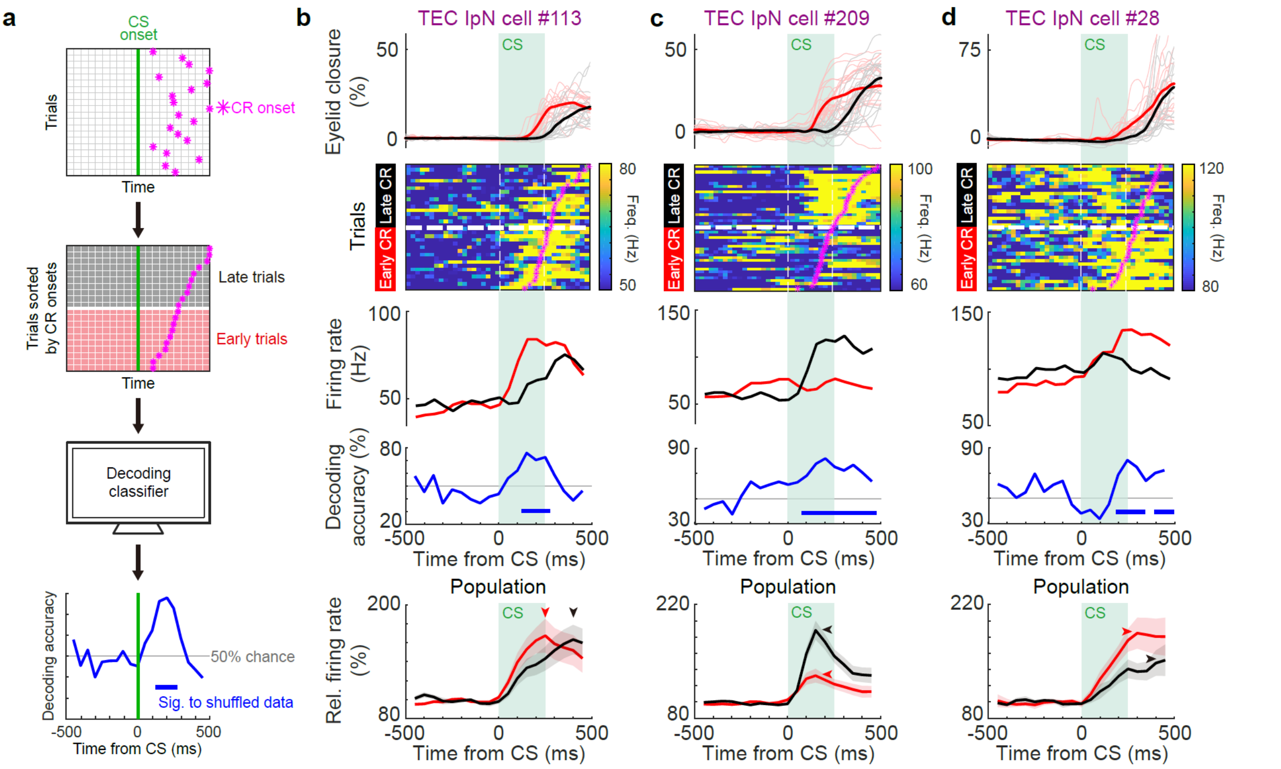


**Extended data Figure 4: Decoding classifier reveals multiplex encoding modes of CR onset timing in IpN neurons. Data related to fig.2.**

**a** Schematics of the decoding classifier pipeline. All trials in one TEC session are sorted based on the CR onsets. The decoding classifier is trained using behavioral and IpN recording data from a subset of trials and applied to the other subset (see details in Methods). **b-d** Multiplex coding strategies of IpN neurons for CR onset timing. Top row: CR traces showing early (red) and late (black) onsets. Second row: instantaneous firing rate of example IpN neurons during early and late CR trials. All trials are sorted based on the CR onsets (magenta dots). Third row: average firing rates of the same example IpN neurons during early and late CR trials. Fourth row: decoding accuracy plotted as the function of time. Grey line indicates 50% decoding accuracy. Significantly decoded epochs are indicated by blue bars. Bottom row: average firing rates of all cells presenting similar coding strategies. Red and black arrowheads indicate the modulation peaks during early and late CRs. **b** IpN neurons encode CR onsets by varying the modulation time (*n* = 11 cells in population); **c, d** IpN neurons encode CR onsets by varying the modulation amplitudes (*n* = 22 and 9 cells in population).

### Extended data Figure 5

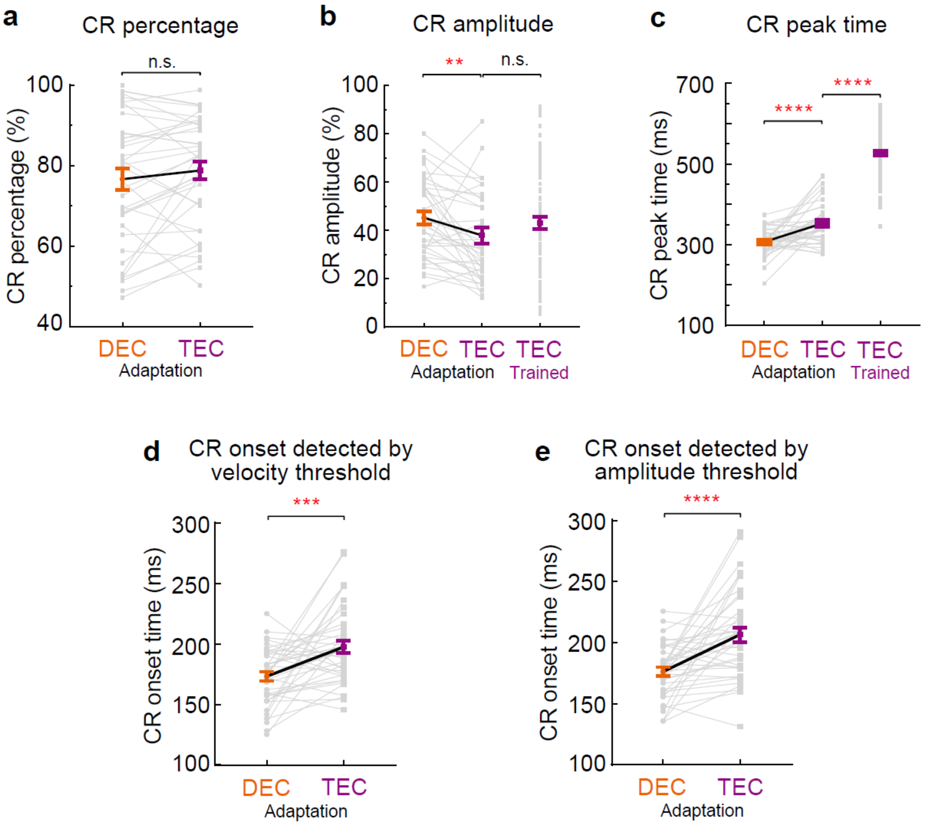


**Extended data Figure 5: Transition of CR kinetics during DEC-to-TEC adaptation. Data related to fig.3.**

**a** CR-trial probability during DEC-to-TEC adaptation ($P = 0.21$, $n = 38$ sessions). **b, c** Comparison of the CR amplitude (**b**) and CR peak time (**c**) in the mice underwent DEC-to-TEC adaptation and the mice trained with TEC during probe trials ($P = 0.0022$ and $P = 0.28$ in **b**, $P < 0.0001$ for both in **c**, $n = 38$ and $n = 92$ sessions). **d, e** The CR onset time during all DEC-to-TEC adaptation recordings which was detected by CR velocity (**d**) and CR amplitude threshold (**e**) ($P = 0.0007$ in **d**, and $P = 1.6 \times 10^{-5}$ in **e**, $n = 38$ sessions for both). Data are shown as mean ± s.e.m., n.s., not significant, $^{**}P \leq 0.01$, $^{***}P \leq 0.001$, and $^{****}P \leq 0.0001$.

### Extended data Figure 6

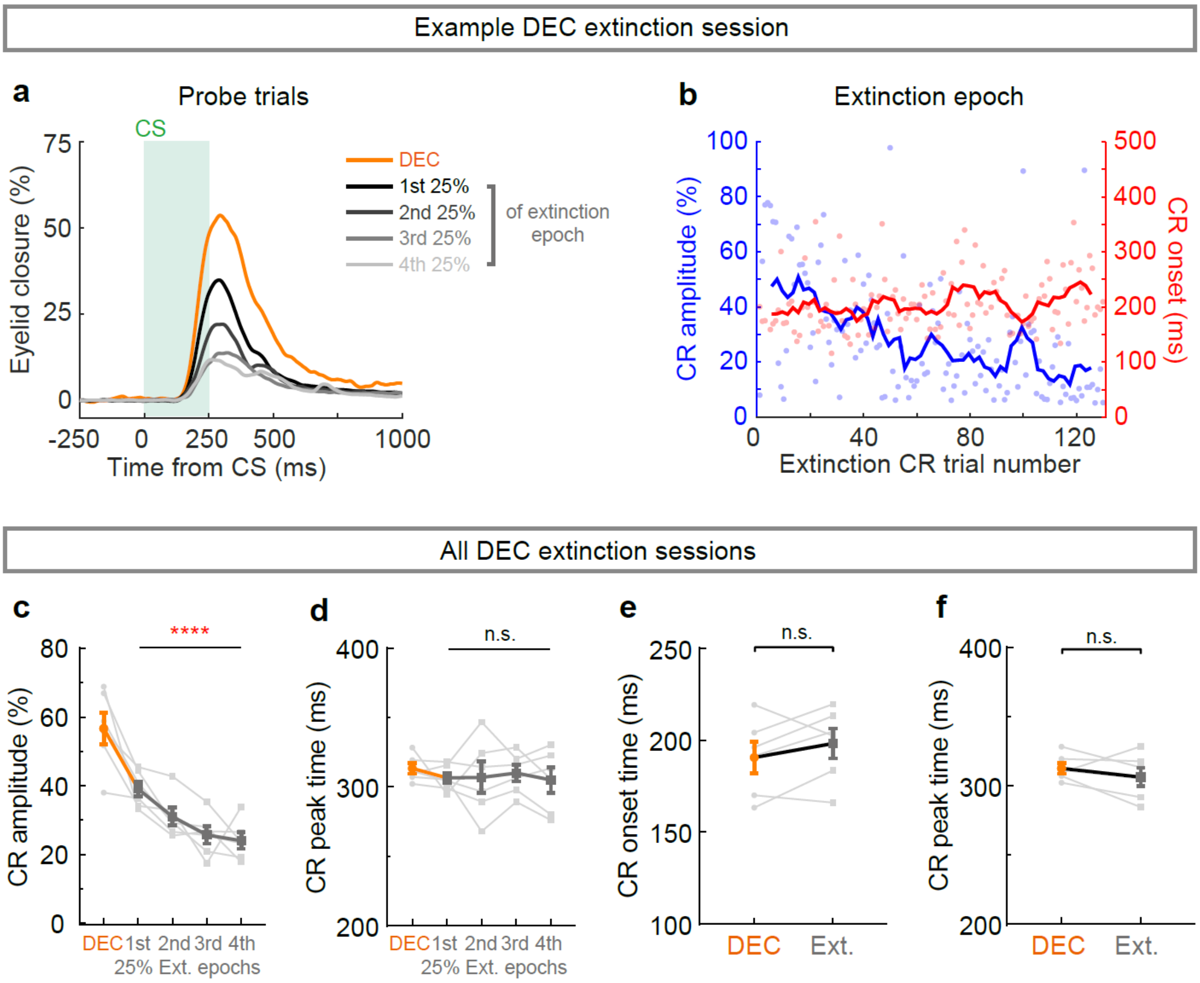


**Extended data Figure 6: The behavior properties of DEC extinction tests. Data related to fig.3.** **a** The average eyelid closure curves of probe trials in DEC epoch (orange) and all four quartiles (black to light grey) in extinction epoch from an example session. **b** The CR amplitude (blue) and CR onset (red) of the example session in (a) in trial-by-trial manner. The solid curves represent to moving average. **c** CR amplitude in the DEC epochs and all four quartiles of extinction epochs ($P < 0.0001$, $n = 6$ sessions). **d** CR peak time in the DEC epochs and all four quartiles of extinction epochs ($P = 0.67$, $n = 6$ sessions). **e, f** CR onset (**e**) and CR peak time (**f**) in the DEC epochs and whole extinction epochs ($P = 0.31$ in **e**, and $P = 0.31$ in **f**, $n = 6$ sessions). Data are shown as mean ± s.e.m., n.s., not significant, and $^{****}P \leq 0.0001$.

**Extended data Figure 7**

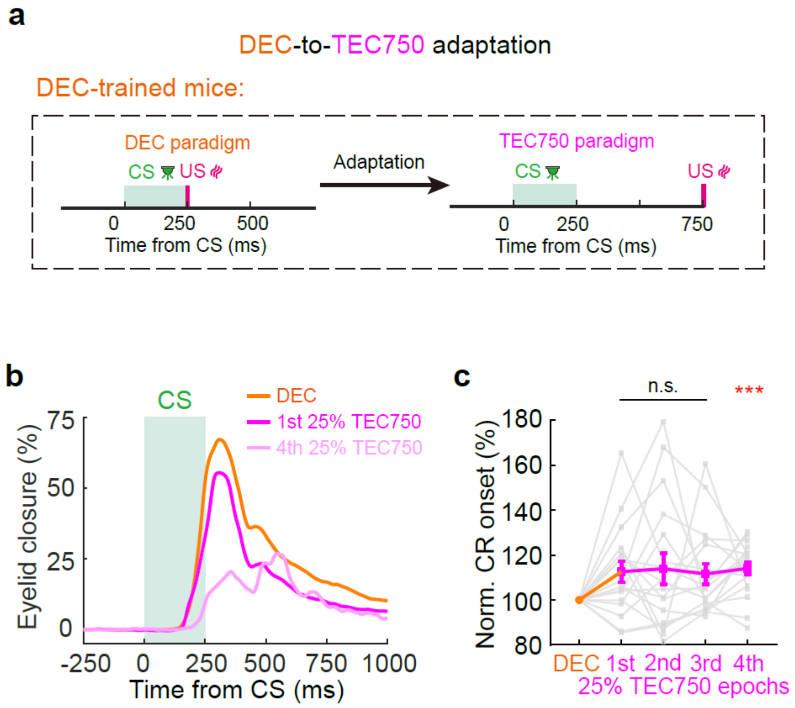


**Extended data Figure 7: The behavior properties of DEC-to-TEC750 adaptation paradigm. Data related to fig.3.**

**a** Experimental procedure for the animal training and paradigm switch to TEC750 in DEC-trained mice. **b** The average probe trial eyelid closure curves of DEC epoch (orange), first (pink) and last (light pink) quarters of TEC epoch from an example session. **c** Normalized CR onset in the DEC epochs and all four quartiles of TEC750 epochs during the adaptation paradigm ($P > 0.05$ for first three quarterlies, and $P = 0.0004$ for the fourth quartiles, $n = 18$ sessions). Data are shown as mean ± s.e.m., n.s., not significant, and $^{***}P \leq 0.001$.

### Extended data Figure 8

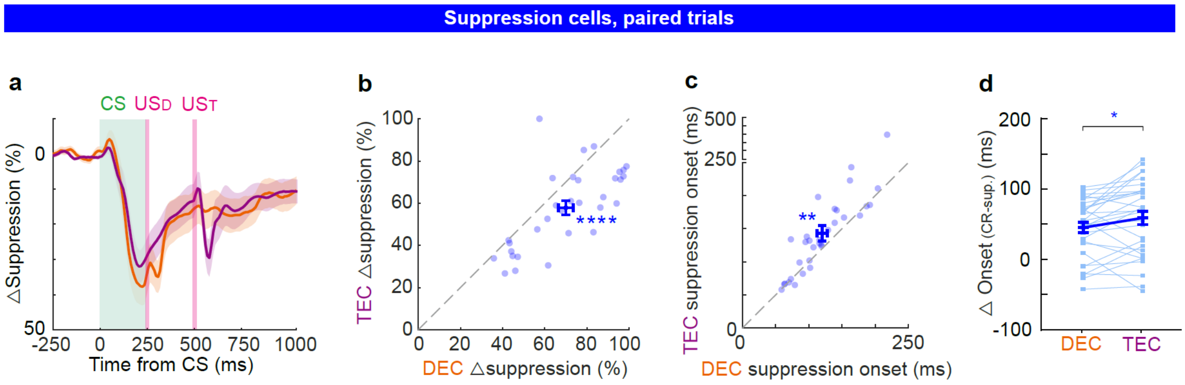


**Extended data Figure 8: DEC-to-TEC switch causes adaptation of the suppression timing and suppression amplitude in a group of IpN neurons. Data related to fig.4.**
**a** Averaged firing rates of suppression cells during DEC (orange) and after adapting to TEC (purple). **b-d** Suppression amplitude (**b**), suppression onset timing (**c**), as well as the interval between suppression timing and CR timing (△onset, **d**) of individual IpN neurons underwent DEC-to-TEC adaptation ($P = 2.54 \times 10^{-5}$ in **b**, $P = 0.0019$ in **c**, and $P = 0.0104$ in **d**, $n = 32$ cells). Data are shown as the mean ± s.e.m., n.s., not significant, *$P \leq 0.05$, **$P \leq 0.01$, and ****$P \leq 0.0001$.

### Extended data Figure 9

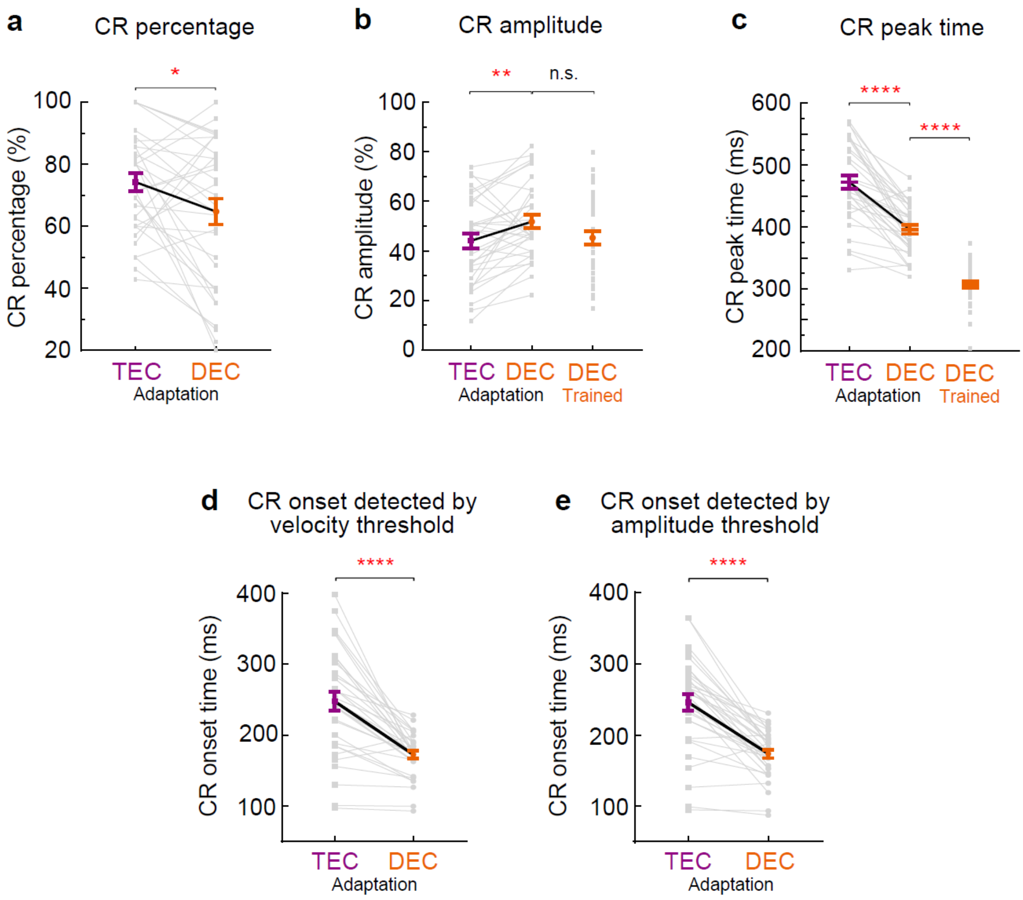


**Extended data Figure 9: Transition of CR kinetics during TEC-to-DEC adaptation. Data related to fig.5.**

**a** CR-trial probability during TEC-to-DEC adaptation ($P$ = 0.02, $n$ = 33 sessions). **b, c** Comparison of the CR amplitude (**b**) and CR peak time (**c**) in the mice underwent TEC-to-DEC adaptation and the mice trained with DEC during probe trials ($P$ = 0.0029 and $P$ = 0.071 in **b**, $P$ < 0.0001 for both in **c**, $n$ = 33 and $n$ = 38 sessions). **d, e** The CR onset time during all TEC-to-DEC adaptation recordings which was detected by CR velocity (**d**) and CR amplitude threshold (**e**) ($P$ < 0.0001, and $n$ = 38 sessions for both). Data are shown as mean ± s.e.m., n.s., not significant, *$P \leq 0.05$, **$P \leq 0.01$, and ****$P \leq 0.0001$.

### Extended data Figure 10

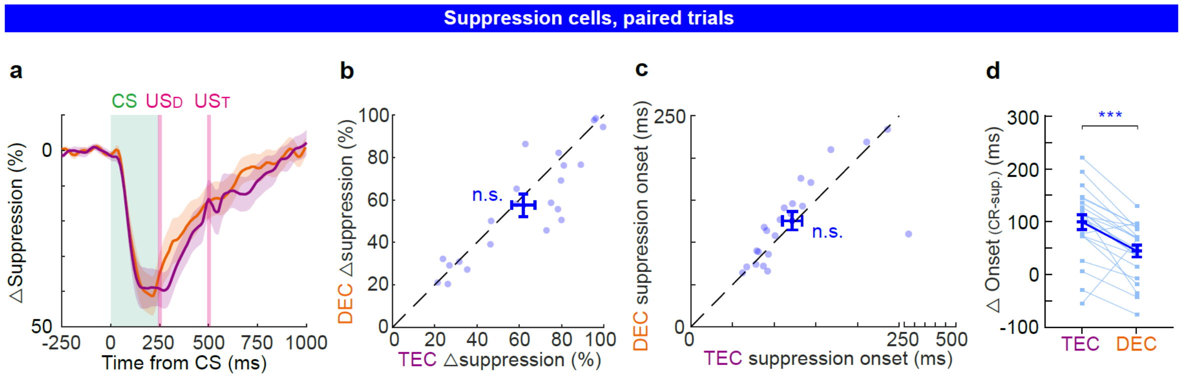


**Extended data Figure 10: Sustained IpN modulation patterns during TEC-to-DEC adaptation. Data related to fig.6.**

**a** Averaged firing rates of suppression IpN cells during TEC (purple) and after adapting to DEC (orange). **b-d** Suppression amplitude (**b**), suppression onset (**c**), and differential onset between suppression and CR (△onset, **d**) of individual neurons during TEC-to-DEC adaptation ($P$ = 0.11 in **b**, $P$ = 0.66 in **c**, and $P$ = 0.0003 in **d**, $n$ = 22 cells). Data are shown as the mean ± s.e.m., n.s., not significant, ***$P \leq 0.001$.

### Extended data Figure 11

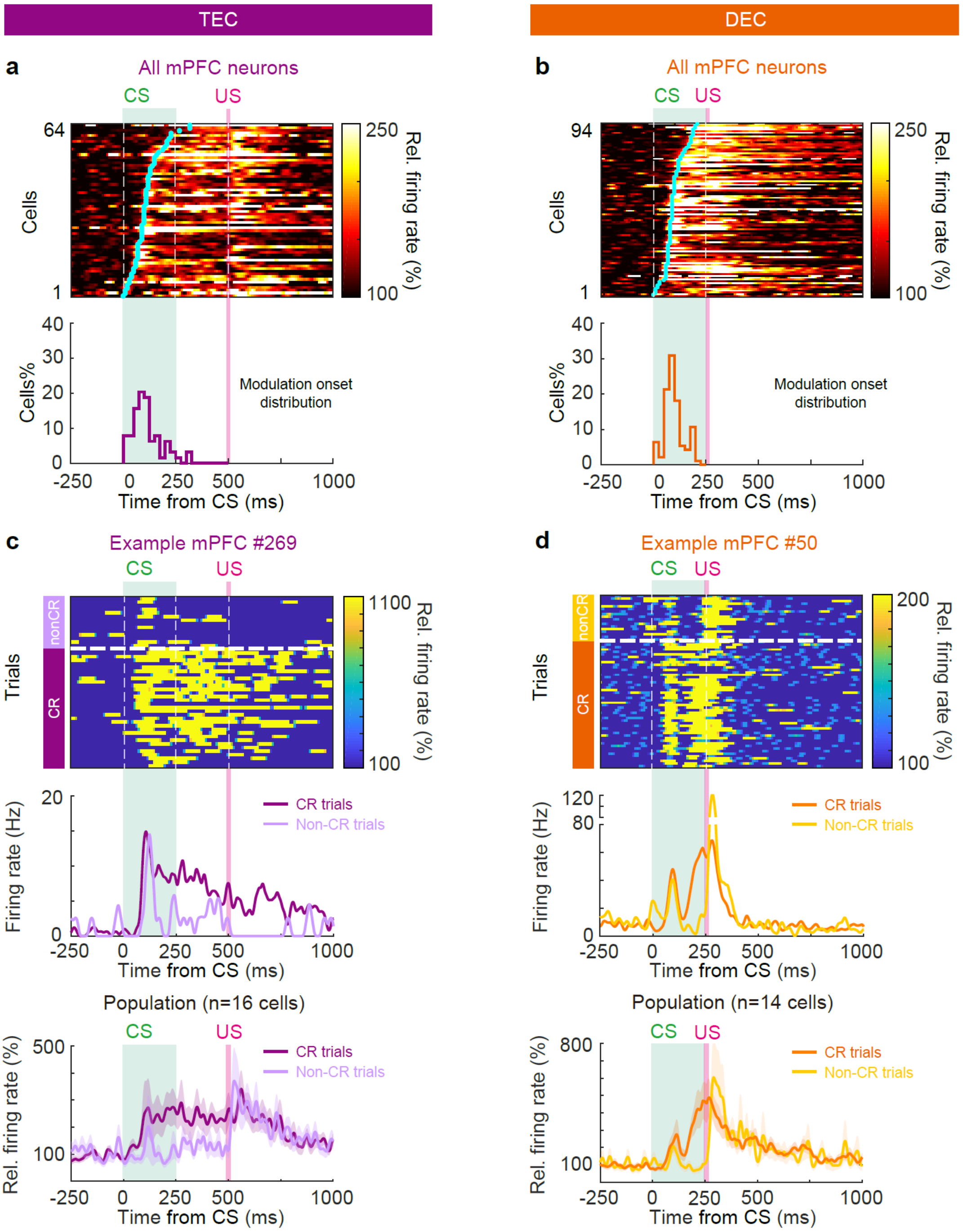


**Extended data Figure 11: Neuronal dynamics of mPFC neurons in TEC and DEC paradigms. Data related to fig.7.**

**a** Top: summary of the mPFC neurons showing task related facilitation during TEC. Each row of the heatmap represents one neuron, and cyan dot represents the facilitation onset. Bottom: distribution of facilitation onsets (*n* = 64 neurons). **b** Same as (**a**), but for the mPFC neuronal activity during DEC (*n* = 94 neurons). **c-d** Comparison of the mPFC activity in CR and non-CR trials during TEC (**c**) and DEC (**d**). Top to bottom: heatmap indicates instantaneous firing rate of an example neuron (top), PSTH of the firing rates of the same example neuron (middle), and (bottom) populational summary for all mPFC neurons during CR and non-CR trials. *n* = 19 cells in TEC recordings and *n* = 14 cells in DEC recordings. Data is shown as the mean ± s.e.m..

### Extended data Figure 12

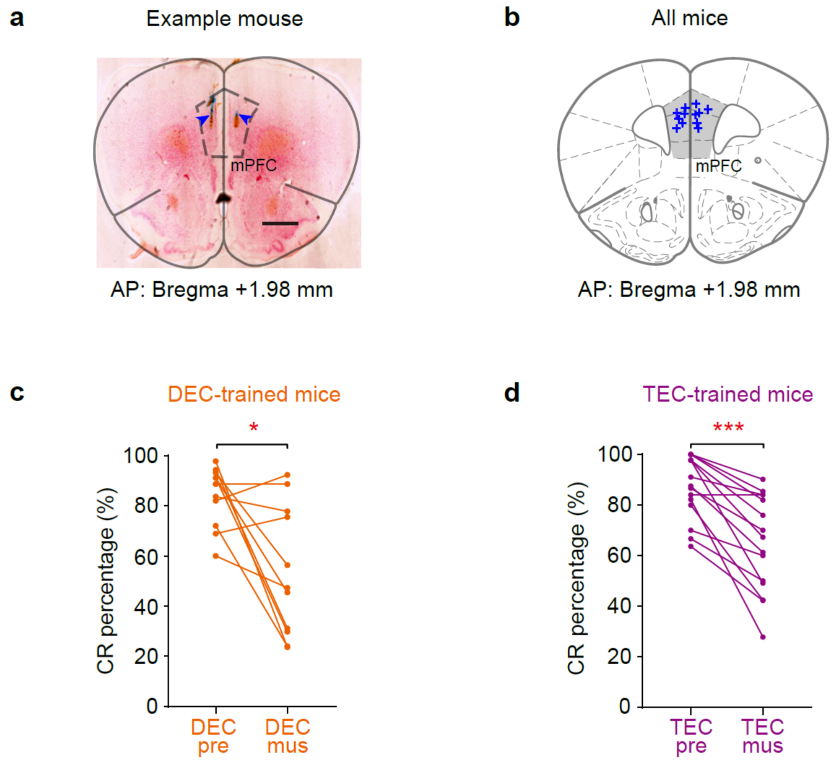


**Extended data Figure 12: Pharmacological inhibition of mPFC impairs CR performance during DEC and TEC. Data related to fig.9.**

**a** Representative histological section showing the sites of bilateral muscimol injection, labeled by alcian blue (arrowheads), in mPFC. **b** Summary of all the muscimol injection sites in 6 mice. **c-d** Comparison of CR-trial probability before and after mPFC inhibition in DEC- (**c**) and TEC-trained (**d**) mice ($P$ = 0.020, $n$ = 11 sessions in **c**, and $P$ = 0.0001, $n$ = 15 sessions in **d**). Data is shown as the mean ± s.e.m., *$P \leq 0.05$, and ***$P \leq 0.001$.
